# Strategies for Efficient Genome Editing Using CRISPR-Cas9

**DOI:** 10.1101/465252

**Authors:** Behnom Farboud, Aaron F. Severson, Barbara J. Meyer

## Abstract

The targetable DNA endonuclease CRISPR-Cas9 has transformed analysis of biological processes by enabling robust genome editing in model and non-model organisms. Although rules directing Cas9 to its target DNA via a guide RNA are straightforward, wide variation occurs in editing efficiency and repair outcomes, both imprecise error-prone repair and precise templated repair. We found that imprecise and precise DNA repair from double-stranded breaks (DSBs) is asymmetric, favoring repair in one direction. Using this knowledge, we designed RNA guides and repair templates that increased the frequency of imprecise insertions and deletions and greatly enhanced precise insertion of point mutations in *Caenorhabditis elegans*. We devised strategies to insert long (10 kb) exogenous sequences or incorporate multiple nucleotide substitutions at considerable distance from DSBs. We expanded the repertoire of co-conversion markers appropriate for diverse nematode species. These selectable markers enable rapid identification of Cas9-edited animals also likely to carry edits in desired targets. Lastly, we explored the timing, location, frequency, sex-dependence, and categories of DSB repair events by developing loci with allele-specific Cas9 targets that can be contributed during mating from either male or hermaphrodite germ cells. Our studies revealed a striking difference in editing efficiency between maternally and paternally contributed genomes. Furthermore, imprecise repair and precise repair from exogenous repair templates occur with high frequency before and after fertilization. Our strategies enhance Cas9 targeting efficiency, lend insight into the timing and mechanisms of DSB repair, and establish guidelines for achieving predictable precise and imprecise repair outcomes with high frequency.

## Introduction

Programmable DNA endonucleases have transformed the analysis of biological processes by enabling targeted genome editing in diverse species (reviewed in (Urnov *et al*. 2010; Joung and Sander 2013; Carroll 2014; Doudna and Charpentier 2014; Chandrasegaran and Carroll 2016). These nucleases catalyze DNA breaks at specified sequences and trigger precise and imprecise repair outcomes via different pathways (Jasin and Haber 2016). Imprecise repair pathways introduce small-to-large insertions and deletions. Precise repair pathways insert specific DNA changes from either homologous chromosomes or exogenous, homologous templates. Our goal is to improve the frequency and fidelity of genome editing by investigating the rules governing repair pathway choices and repair outcomes.

The programmable endonuclease most widely deployed is the *Streptococcus pyogenes* CRISPR-associated protein 9 (*Sp*Cas9 or Cas9) (reviewed in (Mali *et al*. 2013a; Jiang and Doudna 2017). Cas9 is directed to its DNA target by a guide RNA that pairs with 20 bases of target DNA, called the spacer sequence (Mojica *et al*. 2009; Garneau *et al*. 2010; Jinek *et al*. 2012). The chief constraint limiting target choice is the requirement for an NGG motif to border the DNA sequence that is complementary to the spacer. The NGG motif is called the protospacer adjacent motif (PAM). The Cas9-guide RNA ribonucleoprotein (RNP) complex scans the genome, binds to PAM sequences, and melts the adjacent duplex DNA (Sternberg *et al*. 2014). If the neighboring nucleotides are complementary to the guide RNA, Cas9 undergoes a conformational change that activates its nuclease domains to make a DNA double-strand break (DSB) (Anders *et al*. 2014; Sternberg *et al*. 2015).

These basic principles governing Cas9 targeting led to its widespread usage, but the repertoire of strategies that can be used to achieve desired genomic changes has some limitations. In this study, we develop strategies that overcome several impediments to achieve efficient Cas9 targeting and predictable DSB repair outcomes in the nematode *Caenorhabditis elegans*. These approaches can be exploited to improve genome editing across diverse plant and animal species.

We demonstrate that both imprecise and precise DNA repair from a single DSB is asymmetric, favoring repair in only one direction. We have exploited this property to establish guidelines for effective PAM choice and design of single-stranded repair templates to achieve high frequency insertion of desired changes within close proximity (30 bp) to a DSB and on a particular side of the DSB. We also devised efficient strategies to insert long non-homologous fragments of DNA (~ 10 kb) at DSB sites and to engineer small specific changes at considerable distance from a DSB (1.5 kb) or to incorporate a series of nucleotide substitutions throughout an entire locus, all without co-inserting a selectable marker. These strategies are also useful for inserting DNA in sites such as A-T rich regions that are devoid of potential PAMs. These successes required that we optimize Cas9 delivery methods and guide RNA design.

We also expanded the repertoire of Cas9-dependent co-conversion markers to be used in conjunction with the tools to edit targets of choice. These selectable markers, appropriate for diverse nematode species, enable the rapid identification of Cas9-edited animals that are also likely to have desired edits in the targets of choice. This approach is particularly useful when searching for edited targets that fail to cause visible phenotypes.

Finally we devised and exploited an editing strategy to explore the timing, location, frequency, sex-dependence, and categories of DSB repair events. We developed loci with allele-specific targets for Cas9 cleavage that can be contributed from either male or hermaphrodite germ cells during mating. We found that male sperm DNA was generally more permissive to Cas9 editing than hermaphrodite germ cell DNA. The frequency of recovering mutations in the locus of interest is higher if editable alleles of both the target locus and the co-conversion marker are contributed from the same parent during mating. We found that imprecise repair and homology-directed repair from homologous chromosomes or exogenous repair templates occurs at an unexpectedly high frequency after fertilization in embryos.

## Materials and Methods

### Strains

Nematode strains were cultured as described previously (Brenner 1974). N2 Bristol was used as the wild-type *C. elegans* strain and AF16 as the wild-type *C. briggsae* strain.

### Genome editing in self-fertile hermaphrodites versus mated hermaphrodites

Genome editing in *C. elegans* typically involves the delivery of Cas9 and guide RNA components to the gonad syncytium of young adult self-fertile hermaphrodites that have completed sperm production but have ongoing oocyte production. Loci within hundreds of individual meiotic nuclei in each gonad syncytium have the potential to undergo Cas9-dependent cleavage and repair events. This approach has been used for several nematode genome editing studies using Cas9 (Cho *et al*. 2013; Dickinson *et al*. 2013; Friedland *et al*. 2013; Katic and Grosshans 2013; Lo *et al*. 2013; Tzur *et al*. 2013; Waaijers *et al*. 2013; Arribere *et al*. 2014; Kim *et al*. 2014; Paix *et al*. 2014; Zhao *et al*. 2014; Witte *et al*. 2015). Our editing experiments in Figures 1-9 and S1-S4 involved self-fertile hermaphrodites.

**Figure 1.**
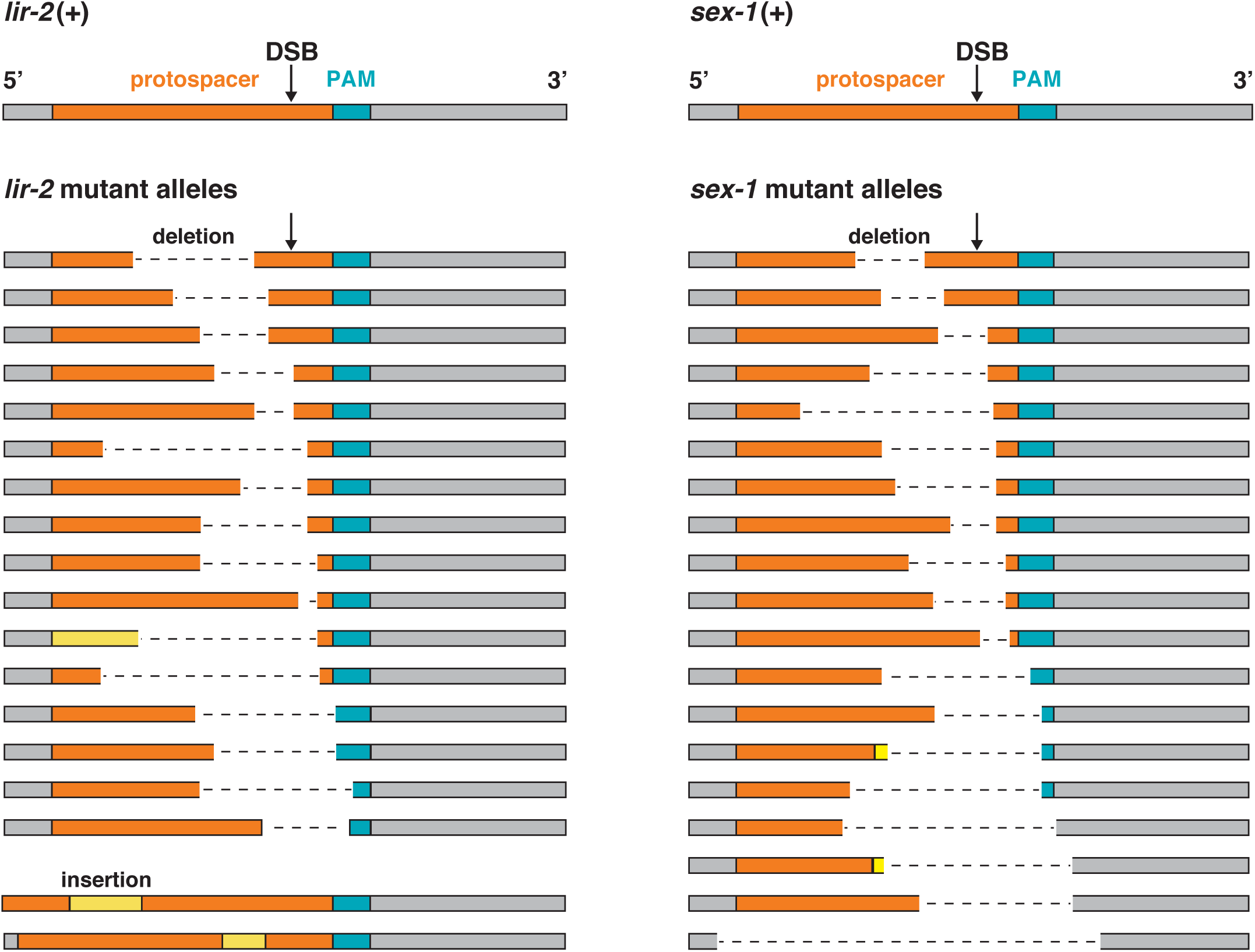
Imprecise repair at Cas9 DSBs causes asymmetric insertion and deletion of DNA sequences in regions 5′ of the PAM. Diagrams depict the protospacer strand from wild-type *lir*-*2* (left) and *sex*-*1* (right) genes, featuring the locations of PAMs (blue), DSBs (arrow), and target-specific sequences (orange). Below are protospacer-strand diagrams showing deletions (dashed lines) and insertions (yellow) caused by imprecise repair at the two genes. All diagrams are drawn to scale. These examples are representative of the *lir*-*2* and *sex*-*1* changes found in edited animals in which *rol*-*6*(gf) was the co-conversion marker. For the last two *lir*-*2* repair examples, only insertions occurred. Because the DNA is drawn to scale and same amount of DNA is shown for all examples, the grey region to the left of the protospacer was shortened to accommodate the length of the insertion. Imprecise repair was biased toward regions located 5′ of the PAM.

For experiments in Figures 10-12 and S5 that explored the timing, location, frequency, sex-dependence, and categories of DSB repair events, we assayed editing in the cross progeny of mated hermaphrodites. We developed loci with allele-specific targets for Cas9 cleavage that could be contributed from either male or hermaphrodite germ cells during mating. Only the maternal or paternal allele of a locus could be edited (as described later in Materials and Methods). We delivered Cas9 RNPs with or without exogenous repair templates to the gonad syncytium of hermaphrodites that had been mated by males for 24 hours. We scored loci of cross progeny hermaphrodites for imprecise repair and homology-directed repair from homologous chromosomes or exogenous repair templates.

### Guide RNAs

For genome editing using sgRNA guides produced from DNA expression vectors (Figures 1, 4B, 7), the relevant target-specific DNA sequences listed in Table S1 were cloned into pRB1017 as described previously (Arribere *et al*. 2014). For editing using Cas9 RNPs, both the target-specific CRISPR RNA guides (crRNA) and the trans-activating crRNA (tracrRNA) were obtained from Dharmacon. All crRNAs, except those used in Figure 7, possessed a 2xMS modification that improves nuclease resistance (Hendel *et al*. 2015; Dowdy 2017). The different editing efficiencies for the *sex*-*1* AG crRNA used in Figure 7 versus the *sex*-*1* AG crRNA used in Figures 10-12 are likely due to the crRNA modification. The efficiencies of all four different crRNAs used in Figure 7 are directly comparable, as are the efficiencies of all crRNAs used in the other figures.

### Repair templates

All ssDNA oligonucleotide repair templates (Table S2) were ordered from Integrated DNA Technologies (IDT) at the 4 nmole Ultramer scale. Single-stranded templates used for examining SNP insertion efficiency 5′ and 3′ of the PAM (Figure 2, A and B, Figure 3, A and B) were PAGE purified to maximize the fraction of full-length templates.

**Figure 2.**
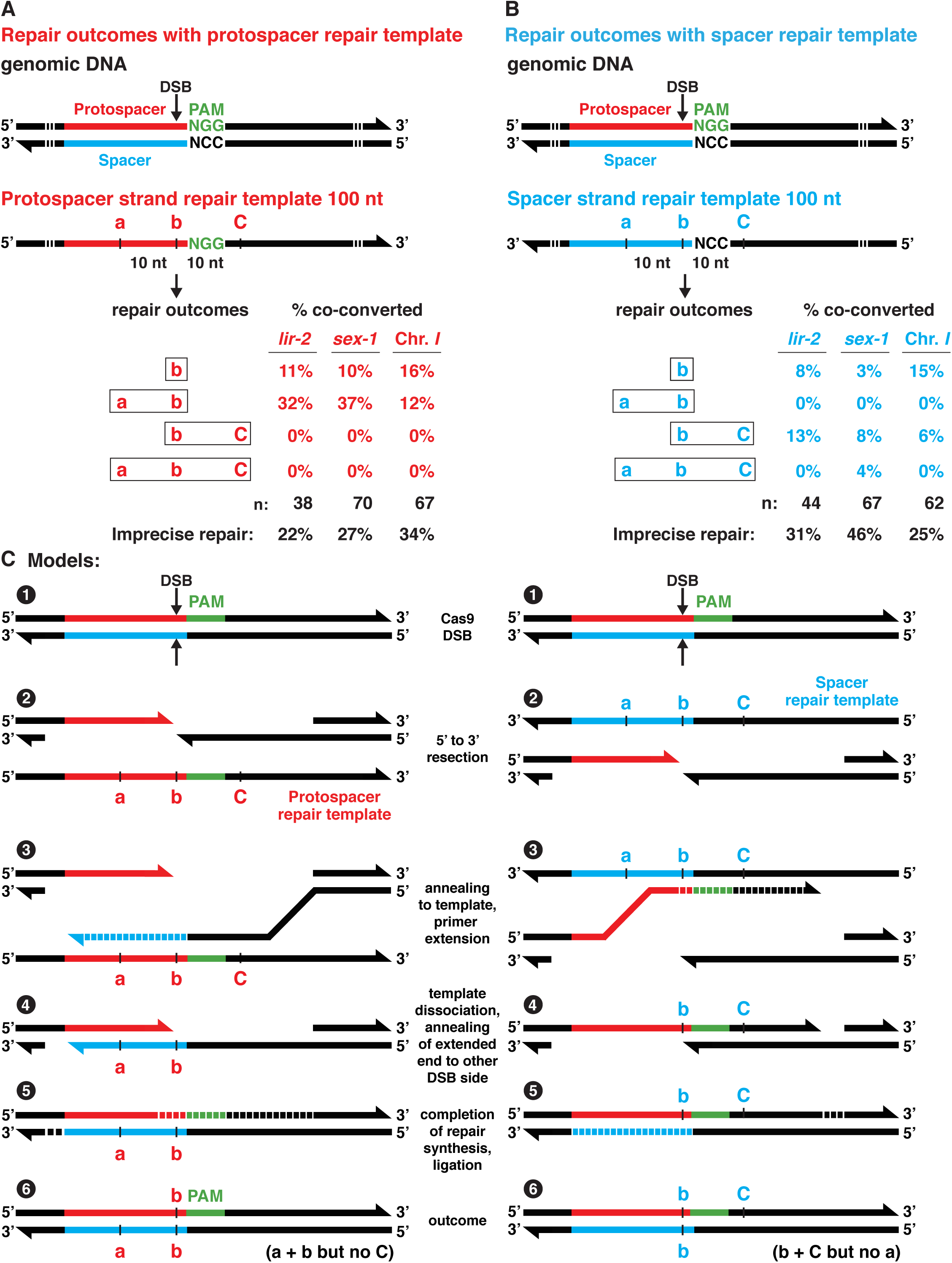
Polarity of homology-directed repair (HDR) from single-stranded templates is dictated by the choice of repair template strand (A,B) Analysis of genome editing experiments at three loci (*lir*-*2*, *sex*-*1*, intergenic Chr I site) comparing HDR outcomes using Cas9 RNPs and single-stranded repair templates of either the protospacer strand (A) or spacer strand (B). Polymorphisms were located in the repair templates at the predicted endogenous DSB sites (SNP b) and at sites both 5′ (SNP a) and 3′ (SNP C) of the PAM (Table S2). Percentages represent the frequencies of diagrammed repair outcomes at each locus relative to the total number of mutants (n) expressing the co-conversion marker in that experiment. Both protospacer and spacer repair templates promoted insertion of SNPs at the DSB site, but the protospacer stand template promoted insertion of SNPs exclusively 5′ of the PAM, while the spacer strand template promoted insertion of SNPs preferentially 3′ of the PAM. Frequency of imprecise repair is also provided for each experiment. For these experiments, *dpy*-*10*(gf) was the co-conversion marker. When precise templated HDR occurred at the *dpy*-*10* locus, the animals exhibited a dominant Rol phenotype and a recessive Dpy phenotype. If imprecise repair occurred at *dpy*-*10*, animals exhibited only a recessive Dpy phenotype. All animals with any of these phenotypes were examined for SNPs. (C) Models show proposed steps in the HDR repair process for protospacer (left) and spacer (right) strand repair templates based on a repair mechanism involving synthesis-dependent strand annealing. Numbers refer to sequential steps.

**Figure 3.**
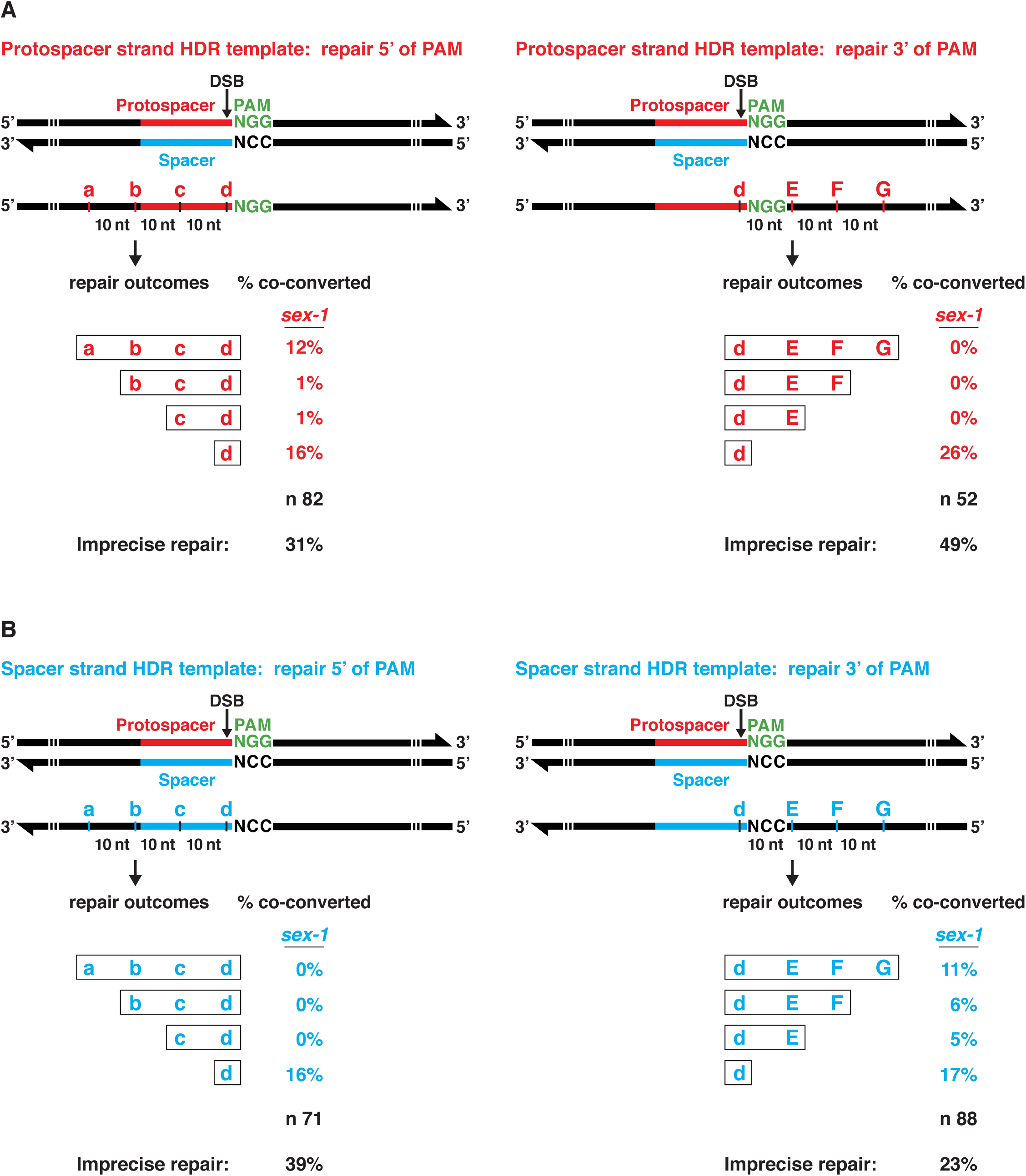
Frequency of SNP insertion from single-stranded repair templates as a function of distance from the DSB (A,B) Shown is the frequency of diagrammed HDR repair outcomes at the *sex*-*1* locus using Cas9 RNPs and either single-stranded protospacer strand (A) or spacer strand (B) repair templates with two SNPs located at the DSB (polymorphism d) and single SNPs in one direction from the DSB, in 10 nt intervals. Insertion of polymorphisms at the DSB was highly efficient regardless of the repair template strand, but protospacer strand templates promoted insertion of SNPs only 5′ of the PAM, while spacer strand templates promoted insertion of SNPs only 3′ of the PAM. Repair outcomes revealed a strong tendency to incorporate all SNPs in one direction from the PAM, not just a subset of SNPs. Repair templates include a 40 nt homology arm, an adjacent 30 nt region with SNPs, a PAM, and a 50 nt homology arm. The percentages of repair outcomes reflect the number of the diagrammed outcomes relative to the total number of *dpy*-*10* mutants (n) in the experiment. Percentage of imprecise repair is also indicated.

Long double-stranded DNA repair templates (Figures 4A, 5, 6, and S2) were purchased as gBlocks from IDT or were made using Gibson assembly reactions to fuse small gBlocks and PCR fragments (Gibson 2011). The templates included silent mutations to eliminate the PAM or to mutate key nucleotides in sequences targeted by guide RNAs. The mutations were designed to prevent Cas9 from cleaving the repair template and the edited genomic locus but not to alter the primary amino acid sequence of the protein encoded by the edited locus. Mating experiments described later in this section confirmed that a single mismatch at the 3′ end of the protospacer can block Cas9 cleavage of the endogenous locus. Sequences for these long dsDNA repair templates are available upon request.

**Figure 4.**
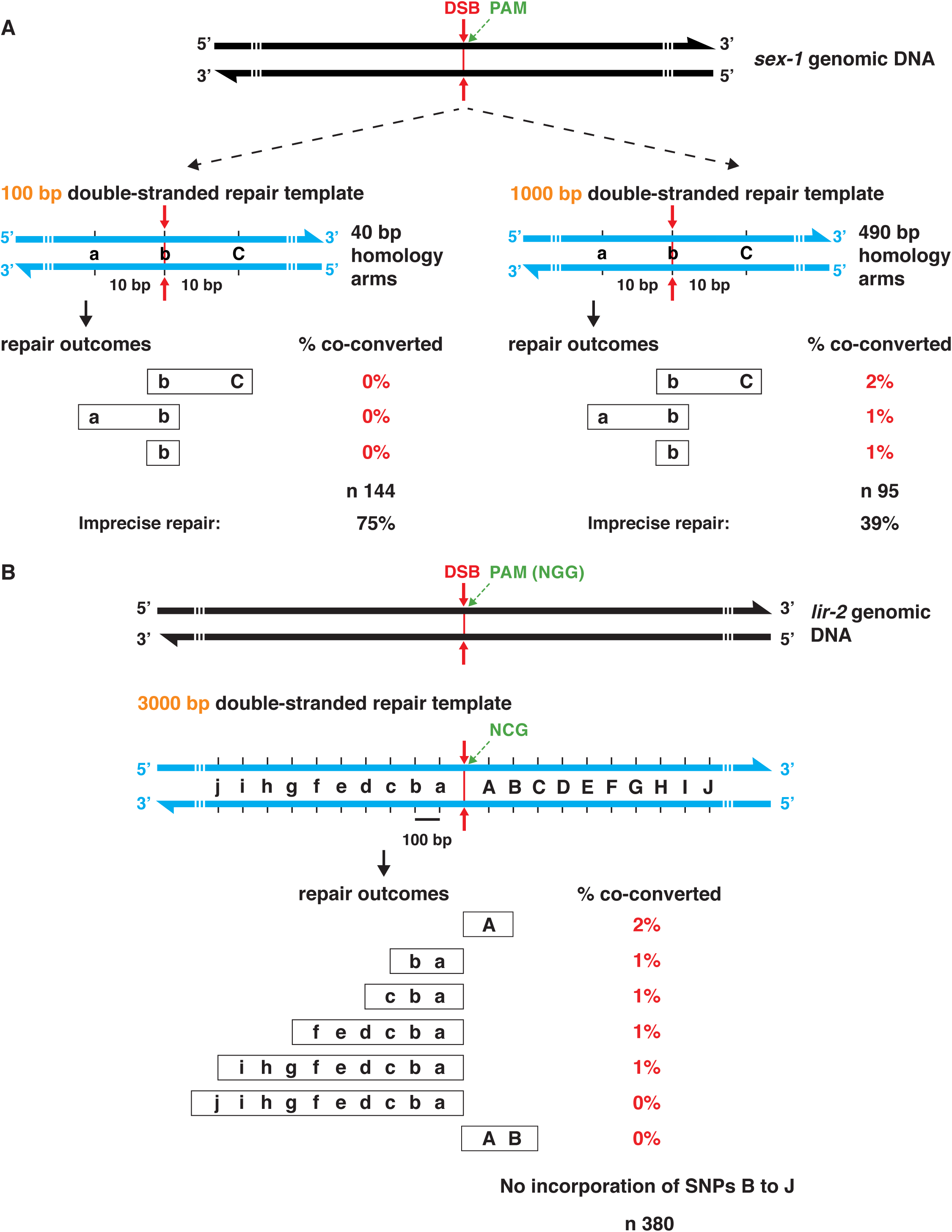
Homology-directed repair from double-stranded repair templates was asymmetric relative to the DSB and less efficient than repair from single-stranded templates (A left) The experiment in Figure 2A was reproduced in design except for the use of a linear 100 bp double-stranded repair template instead of a 100 nt single-stranded repair template. Unexpectedly, no HDR was observed, although 75% of the *dpy*-*10* mutants had undergone imprecise repair at the DSB in *sex*-*1*. (A right) The experiment in the left panel was repeated, except the linear double-stranded repair template (1 kb) had 490 bp homology arms instead of 40 bp arms. The outcome was more successful, but the frequency of HDR was greatly reduced compared to the frequency of HDR in Figure 2A with single-stranded repair templates. (B) Analysis of SNP insertions relative to the distance from the DSB. The 4 kb double-stranded repair template had 10 polymorphisms at ~ 100 bp intervals on both sides of the DSB and 500 bp homology arms. Each of the 20 polymorphisms created a *Hind*III restriction site for analyzing repair outcomes. Precise HDR repair was infrequent and exhibited a striking directionality in which infrequent insertions occurred up to 900 bp from the DSB, but predominantly only 5′ of the PAM. For (A,B), the number of animals with the diagrammed insertion was expressed as the percentage of the total number of*dpy*-*10* Dpy or Rol mutants (n). Cas9 RNPs were used in (A), and DNA expression vectors were used for Cas9 and the guide RNAs in (B). The 4 kb repair template was introduced as plasmid DNA.

**Figure 5.**
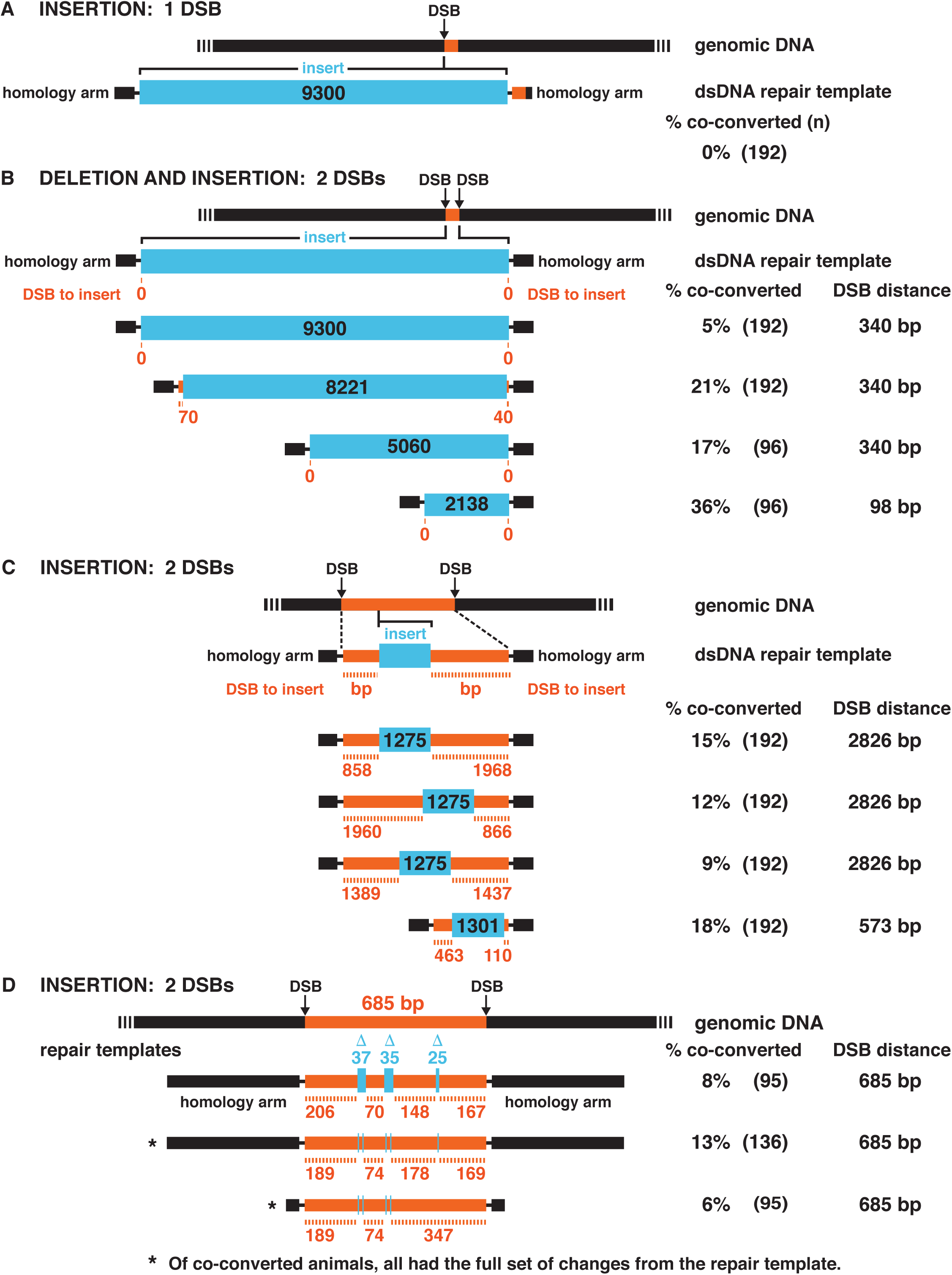
Two neighboring DSBs promote insertion of large DNA fragments (A) Insertion of a 9300 bp reporter transgene was not successful in experiments involving a single Cas9 DSB and a double-stranded repair template with 500 bp homology arms (black and orangeblack) flanking the DSB. See also results in Figure S2. (B) Insertion of long DNA fragments (blue) was efficient in *dpy*-*10* co-conversion experiments when two neighboring DSBs were made using two different guide RNAs. In this experimental design, the DNA between the DSBs (340 bp or 98 bp) was deleted when the large DNA fragment was inserted. Double-stranded repair templates with 500 bp homology arms (black) were injected as plasmid DNA along with the Cas9 RNPs. For insertion of the 8221 bp reporter, the repair template included the standard 500 bp homology arms and additional 70 bp or 40 bp regions of homology to the DNA between the DSBs, making the total length of deleted DNA be 230 bp not 340 bp. For all experiments in this figure, the term “DSB to insert” refers to the distance between the DSB and the novel DNA to be inserted. PAM orientation was IN/IN for the top 3 examples and IN/OUT for the bottom example (see Figure 6 legend for the orientation key). (C) Insertion of long DNA fragments at wide-ranging distances from both DSBs was accomplished by a variation in the double-stranded repair template. The template included all the DNA between the DSBs (orange) and positioned the new DNA to be inserted (blue) at different locations within the DNA spanning the DSBs. In this design, no DNA was deleted at the endogenous locus; only new DNA sequences were inserted. DSB locations for the last example differed from those of the other three examples. PAM orientation was OUT/OUT for the top three examples and OUT/IN for the bottom example. (D) Using a similar design of double-stranded repair template as in (C), genome editing was accomplished at multiple sites in a large region of genomic DNA. In the top example, small deletions (blue) were introduced along the DNA, while in the bottom example, 3 bp substitutions (blue) were inserted at five different locations. PAM orientation was IN/IN for the 3 examples. For (A-D), the number of animals with the diagrammed insertion was expressed as the percentage of total *dpy*-*10* Rol or Dpy mutants given in parenthesis. Homology arms for all repair templates were 500 bp, except for the bottom example in D, which had 50 bp homology arms. All repair templates for (A) except the 3128 bp template were injected as purified plasmid DNA along with the Cas9 RNPs. Repair templates in (C, D) were linear double-stranded DNAs. All diagrams were drawn to scale.

**Figure 6.**
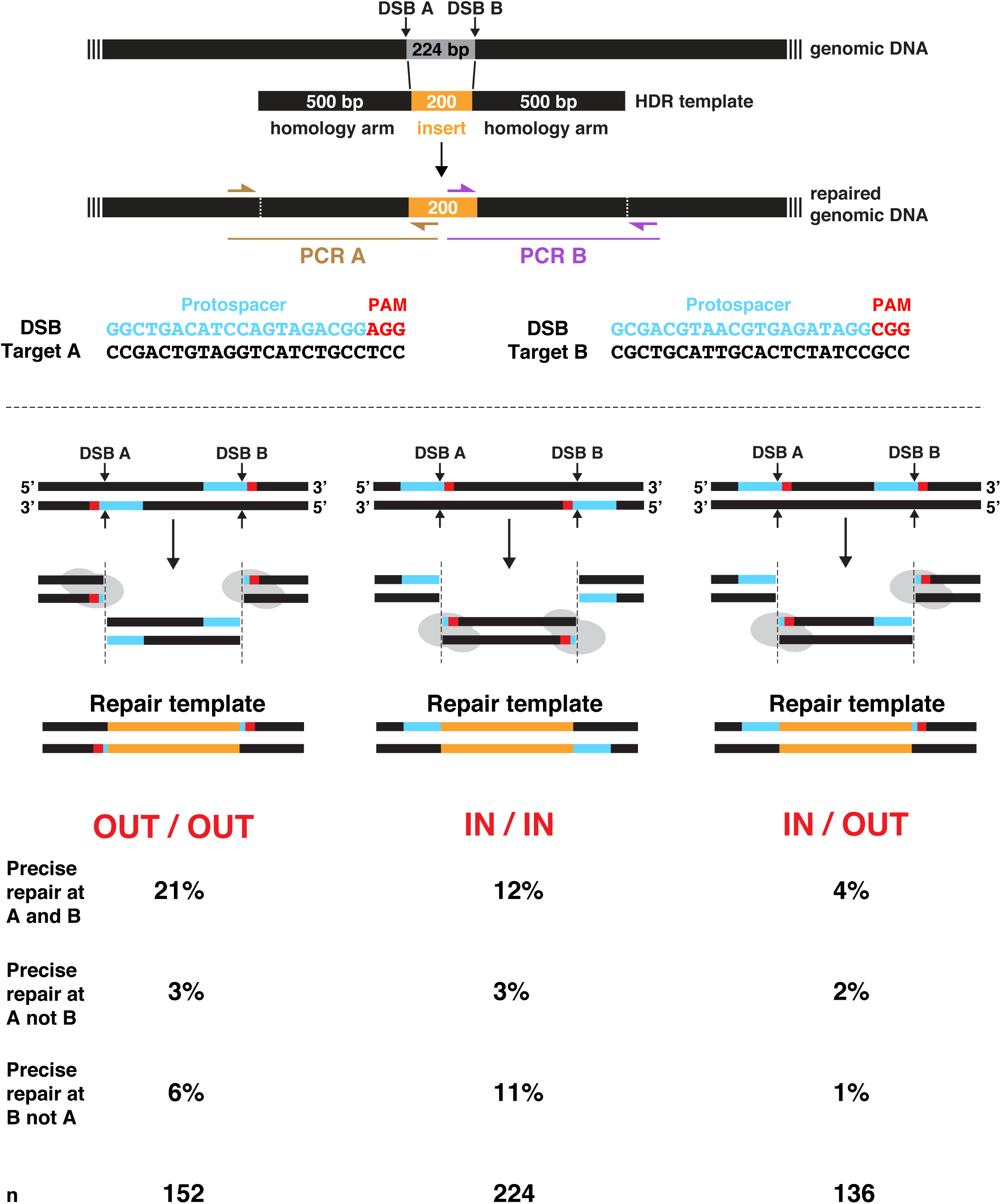
Orientation of PAMs at two nearby Cas9 cleavage sites influences HDR efficiency Cas9 was directed simultaneously to two locations in close proximity (224 bp, Target A and Target B) to make two DSBs and thereby stimulate insertion of an exogenously supplied homologous repair template. The effect of relative PAM orientation on HDR efficiency was assessed by analyzing three strains that had identical sequences at the insertion sites except for the orientation of Targets A and B. The PAM IN orientation placed the PAM within the DNA sequence excised from the chromosome. The PAM OUT orientation left the PAM on the chromosome end after DNA excision. The double-stranded DNA repair templates had 500 bp homology arms corresponding to genomic sequences adjacent to DSB A and DSB B. The three repair templates used for the three different strains were identical except for the cleavage remnants of Targets A and B included in the template, as shown in the diagram. All three templates included the same novel DNA to be inserted (orange). Precise HDR resulted in replacement of the 224 bp region (gray) between DSB A and DSB B with 200 bp of exogenous DNA sequence from the repair template (orange). Only the PAM (red) and protospacer (blue) are shown on the diagrams of target DNA and repair templates. Sanger sequencing of PCR products from the brown primer pair for Target A and the purple pair for Target B was used to screen co-converted Dpy or Rol animals for precise repair at A and/or B (see Methods). Percent (%) co-converted F1s with precise or imprecise repair at Cas9 targets is shown for each PAM orientation. (n) is the total number of co-converted F1s scored by PCR. The percent of repair from the OUT/OUT orientation was significantly greater than that for the IN/IN orientation (*P* = 0.046, chi-square) or the IN/OUT orientation (*P* < 10^−4^, chi-square). The percent of repair from the IN/IN orientation was significantly greater than that for the IN/OUT orientation (*P* = 0.007, chi-square).

The double-stranded repair template *pGEM7z*-*lir*-*2* was constructed to examine the relationship between distance from the Cas9 cleavage site and efficiency of homology-directed repair from an exogenous template (Figure 4B). A variant of the 3 kb region centered on the crispr_bf8 (Table S1) target site was cloned into pBluescript KS(+) using Gibson assembly of overlapping gBlocks (Gibson 2011). The template carried an altered PAM (GGG to GGA) to prevent cleavage of the template and the edited endogenous locus. On each side of the altered PAM, ten *Hind*III sites were created at ~ 100 bp intervals by introducing mutations that changed 1-4 bp. Approximately 500 bp of uninterrupted *lir*-*2* homologous sequence flanked the core 2 kb region with the new *Hind*III sites.

### Genome editing using DNA vectors to express Cas9 and guide RNAs

For editing involving DNA plasmids to express Cas9 and sgRNA guides, injection mixes included 25 ng/μl pRB1017-derived sgRNA plasmid (for each sgRNA), 50 ng/μl pDD162 Cas9 plasmid, and 500 nM *dpy*-*10*(*gf*) or *rol*-*6*(*gf*) single-stranded oligonucleotide repair template. The double-stranded pGEM7z-*lir*-2 repair template was used at 300 ng/μl (Figure 4B). All plasmids were purified using Qiagen’s Midi Plasmid Purification Kit. Injected P0 worms were allowed to recover for 2-3 hr at 20° before being transferred to 25°. After three days, Rol and/or Dpy worms were picked to individual plates, allowed to produce self-progeny, and screened for mutations at loci of interest.

### Genome editing using Cas9 ribonucleoprotein complexes

Cas9 RNPs were prepared for injection as described previously (Paix *et al*. 2015). Briefly, 20 μl injection mixes consisting of 8.75 μM Na-HEPES pH 7.5, 115 μM KCl, 15 μM Cas9-NLS (QB3-UC Berkeley Core facility), 15 μM *dpy*-*10* crRNA guide (crispr_bf32), 15 μM for each crRNA guide to a locus of interest, 42 μM tracrRNA, and 500 nM *dpy*-*10* single-stranded oligonucleotide repair template were assembled on ice. Any additional ssDNA or dsDNA repair templates were used at a final concentration of 500 nM or 350 ng/μl, respectively. RNP cocktails were made and delivered by pipetting all reagents into a tube on ice, incubating at 37° for 15 min, centrifuging at 13,000xg for 1 min, loading into a microinjection needle, and injecting into gonads of young adult hermaphrodites. Injected P0 hermaphrodites were treated as described above.

### Co-conversion strategy using ben-1 in C. elegans and C. briggsae

We developed *ben*-*1* as a co-conversion marker for both *C. elegans* and *C. briggsae* (Figure 8, A and B). *ben*-*1* encodes β-tubulin, and *ben*-*1* mutations confer resistance to benzimidazole (benomyl), which binds to β-tubulin, inhibits microtubule polymerization, and induces paralysis, uncoordinated (Unc) movement, and a dumpy (Dpy) morphology in both strains (Chalfie and Thomson 1982; Driscoll *et al*. 1989).

To establish the benzimidazole concentrations required to induce the desired phenotypes in *C. elegans* and *C. briggsae*, we placed young adult N2 or AF16 worms on NG agar plates containing 5 μM, 7.5 μM, 10 μM, 12.5 μM, or 15 μM benzimidazole, incubated them at 25° for 3 days, and examined their F1 progeny. Growth on 7.5 μM benzimidazole caused the expected phenotypes in *C. elegans*, while growh on 12 μM benzimidazole was required in *C. briggsae*.

The Unc and Dpy phenotypes resulting from benzimidazole treatment were efficiently suppressed by loss-of-function *ben*-*1* mutations induced by injecting Cas9 RNPs mixtures that included the crispr_bf41 guide and BF-2035 repair template in *C. elegans* and the crispr_bf39 guide and BF-2036 repair template in *C. briggsae* (Tables S1 and S2). Both single-stranded repair templates introduced a premature in-frame stop codon at the homologous locus to knock out gene function. To simplify detection of precise edits, repair templates were designed to introduce a new *XbaI* site into the *C. elegans ben*-*1* locus and a new *NdeI* site into the *C. briggsae* locus.

Worms injected with repair templates and Cas9 RNPs targeting *ben*-*1* and a locus of interest were allowed to recover for 2-3 hr at 20° and then transferred to 25° for 2-3 days. Mobile non-Unc non-Dpy animals (produced from successful *ben*-*1* editing) were picked to individual plates, allowed to produce self-progeny at 25°, and screened for edits at loci of interest. Growth at 25° allows detection of both heterozygous and homozygous *ben*-*1* mutations, which are semi-dominant or dominant at 25° but recessive at 15°.

### Co-conversion selection using zen-4 in C. elegans

We developed *zen*-*4*(+) as a co-conversion marker to enable the selection of fully wild-type progeny produced by *zen*-*4*(*cle10*ts) lethal mutants injected with Cas9 RNPs and repair templates (Figure 9, A and B). The *zen*-*4*(*cle10*ts) mutation causes embryonic lethality within 30 min of upshift from the 15° permissive temperature to the 25° non-permissive temperature. The selectable *zen*-*4*(+) repair event restores both DNA and protein sequences to wild-type sequences.

For co-conversion using *zen*-*4*(+) repair templates, young *zen*-*4(cle10*ts) adult hermaphrodites grown at 15° were injected quickly at 20° with a mixture containing Cas9 RNPs with *zen*-*4* guide RNAs (crispr_bf46; Table S1) and the wild-type repair template (BF-2046; Table S2) to convert the three *zen*-*4(cle10*ts) SNPs to fully wild-type sequences (Figure 9). After injection, worms were allowed to recover on NG agar plates at 15° for 24 hr. After 24 hr, the injected P0 worms were transferred to new NG agar plates, and both sets of plates (those with the injected P0s and those with F1 embryos laid for 24 hr) were transferred to 25° for three days. Both *zen*-*4*(+) / *zen*-*4(cle10*ts) and *zen*-*4*(+) / *zen*-*4*(+) embryos hatched and developed into phenotypically normal, fertile adults. In contrast, *zen*-*4(cle10*ts) homozygous mutant animals that hatched prior to the 25° shift developed to adulthood but were Unc, scrawny, and sterile, making them clearly distinguishable from wild-type worms. Mutant embryos that failed to hatch prior to the 25° temperature shift were dead after the shift. Confirmation for the conversion of one or both of the *zen*-*4(cle10*ts) alleles to *zen*-*4*(+) was achieved by digesting a PCR-amplified *zen*-*4* amplicon with *AluI* (Figure 9A). The *zen*-*4(cle10*ts) amplicons were cleaved, but the *zen*-*4*(+) amplicons remained intact. Loci of interest were examined for edits in the *zen*-*4*(+) worms.

### Screening strategies to assess editing at loci of interest

In all experiments presented, co-conversion protocols were used to enrich for worms with genome editing events at loci of interest (Arribere *et al*. 2014; Kim *et al*. 2014; Ward 2015). P0 animals injected with reagents to edit a co-conversion marker and a locus of interest were allowed to lay progeny for two to three days, and F1 progeny expressing the phenotype of a co-conversion marker, *rol*-*6*(gf), *dpy*-*10*(gf), *ben*-*1*(lf) or *zen*-*4*(+), were screened for edits at loci of interest, either after laying progeny or without laying progeny, according to the experimental design.

When using *dpy*-*10* as a co-conversion marker, we found, as expected, that precise repair of one allele with an exogenous repair template that introduces the nucleotide changes found in *dpy*-*10(cn64)* recapitulates the dominant left-hand roller (Rol) phenotype of *dpy*-*10(cn64)* / + animals. However, imprecise repair of a single allele of *dpy*-*10* or heteroallelic combinations of different *dpy*-*10* lesions can yield a variety of Rol, Dpy, or Dpy-Rol phenotypes (Levy *et al*. 1993). We examined animals exhibiting that spectrum of *dpy*-*10* phenotypes for mutations in our gene of interest.

For Figure 1, after individual F1 *rol*-*6*(gf) animals laid F2 progeny on agar plates for 2-3 days, they were transferred to separate wells of a 96-well plate and then lysed using protocols described previously (Farboud and Meyer 2015). Locus-specific PCR and Sanger sequencing of PCR products were performed on DNA from lysed F1s using oligonucleotides in Table S3. Imprecise repair was detected by either the presence of overlapping traces beginning near the DSB site (heterozygous imprecise edits) or single mutant traces (homozygous imprecise edits). F2 progeny from F1 worms carrying heterozygous imprecise repair events were cloned, and their loci of interest were amplified by PCR and then assayed by Sanger sequencing to identify F2 homozygous mutants and determine the exact sequence changes caused by imprecise repair.

To assay genome editing events in experiments of Figures 2, 3, 4A, 7, 10-12, and S5 *dpy*-*10* mutant F1 progeny were lysed in individual wells of a 96-well plate. Loci of interest were PCR amplified using primers in Table S3 and analyzed by Sanger sequencing. For Figures 2, 3, 4A, 7, 10-12, and S5 sequencing chromatograms revealed whether loci had heterozygous programmed HDR events (double traces at the site of SNP insertion), homozygous HDR events (single mutant trace), or no editing events (single wild-type trace). For all five figures, homozygous and heterozygous imprecise repair was detected as in Figure1, except clonal analysis of F2 worms was not used to identify indel endpoints.

**Figure 7.**
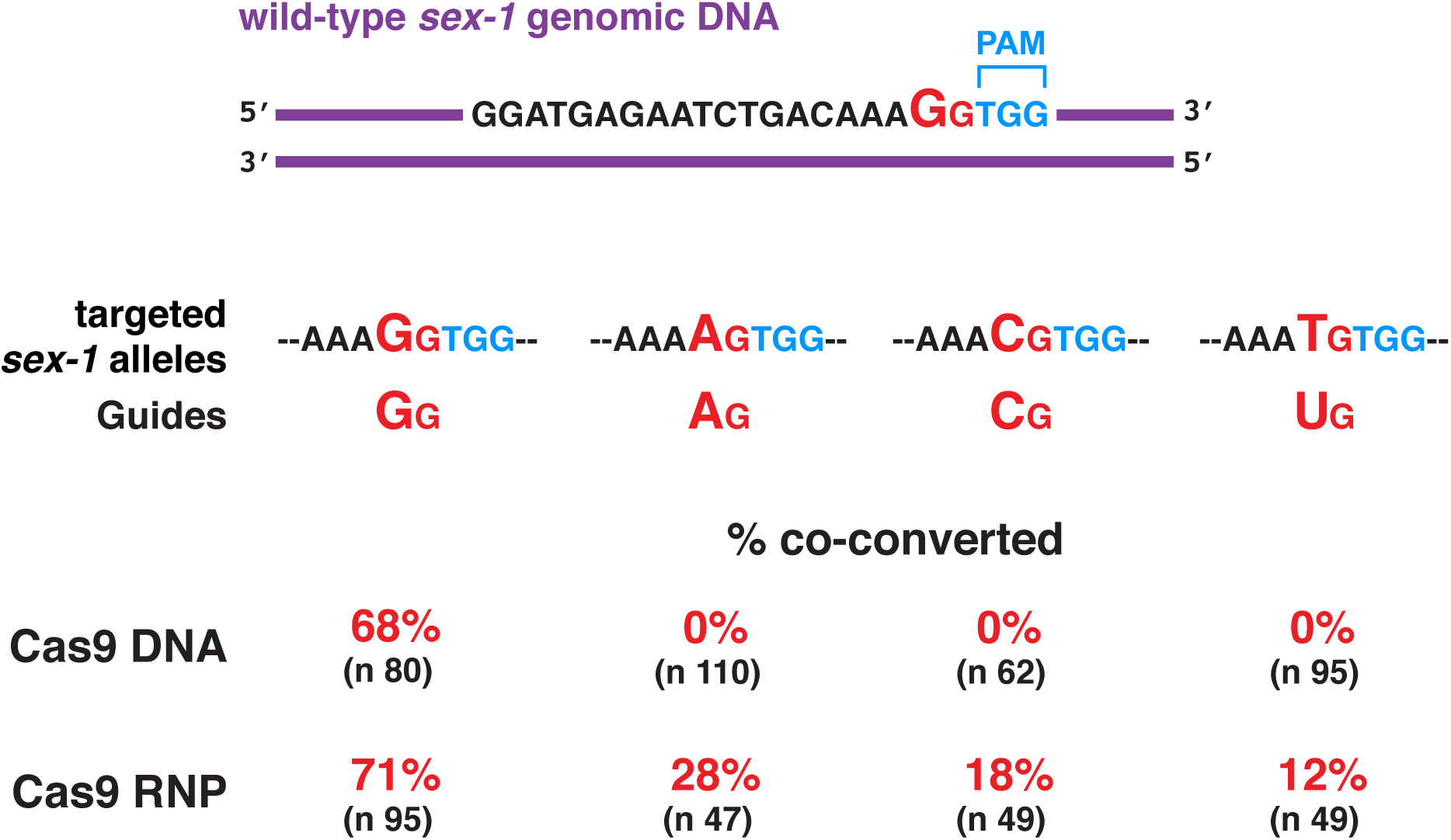
Optimization of guide RNA design and Cas9 delivery method Imprecise repair outcomes in isogenic *sex*-*1* strains differing only by a single base pair change enabled comparison of the effectiveness of four RNA guides differing only in the penultimate nucleotide at the 3′ end of the target-specific sequence. Effectiveness of two different Cas9/guide delivery methods was compared for each guide: RNPs versus DNA expression vectors. Using the DNA expression vector delivery approach, only the GG guide induced imprecise repair. Cas9 RNPs were more permissive, but the GG guide was still more effective than the other guides. Percentages represent the frequencies of imprecise repair events relative to the total number of *dpy*-*10* co-converted F1s (n).

To determine the length of repair tracts templated from dsDNA repair templates harboring *Hind*III sites at 100 bp intervals (Figure 4B), 1.6 kb fragments were amplified on the 3′ and the 5′ side of the PAM using primers listed in Table S3. Amplicons were digested with *Hind*III, and the restriction fragments resolved through electrophoresis on a 2.5% agarose gel. The banding pattern revealed the distance of the terminal-most SNP from the Cas9 DSB. If the entire repair template had been inserted, the amplicon would be reduced to ten ~100bp fragments and the ~ 600 bp fragment from the homology arm that lacks *Hind*III sites. If partial insertion occurred, the size of the largest fragment would reveal the length of DNA lacking *Hind*III sites. Subtraction of the largest fragment length from the full 1.6 kb amplicon length revealed the approximate distance from the DSB that the HDR insertion stopped.

To monitor the insertion of large DNA fragments (Figures 5 and S2), individual F1 *dpy*-*10* mutant worms that laid F2 progeny on individual agar plates were transferred to separate wells of a 96-well plate and lysed. For Figures 5A-C and S2, PCR was performed using primers that yielded product only when the desired genomic insert was present. One primer annealed to the inserted sequence, while the other primer annealed to adjacent chromosomal sequences absent from the repair template. This approach prevented false-positive amplification of repair templates present in extrachromosomal DNA arrays. A second PCR reaction using primers that bound just outside the two Cas9 target sites yielded product only from chromosomes lacking large insertions.

The F2 progeny of F1 animals that tested positive for the insertion were also cloned, allowed to lay F3 progeny, and then genotyped to identify animals that were homozygous for the desired insert. Homozygotes produced insert-specific PCR amplicons, but no PCR amplicons from primers specific for DNA lacking the insert. The accuracy of insertion was confirmed by Sanger sequencing across the entire inserted genomic region of homozygotes from at least 4 independent insertion events. Insertions were precise, with no sequence changes near the DSBs.

For Figure 5D, PCR was performed on DNA from F1s to amplify across the genomic insertion site, with one primer annealing to genomic sequences absent from the repair template. Sanger sequencing was then performed to assess the presence of desired small deletions and point mutations. F2 progeny of F1s that were heterozygous for the desired mutations were cloned and retested by Sanger sequencing to identify homozygous mutants. The full set of desired sequence changes was present in each independent insertion event (8 insertions for the deletions and 18 for the point mutations).

For Figure 6, two sets of PCR reactions were performed on lysed F1s from each strain to test for insertions at DSB A and DSB B. For each reaction, one primer annealed to genomic sequence located outside the bounds of the repair template and a second primer annealed to novel DNA within the repair template. Precise repair at A and B was indicated by properly sized PCR A and PCR B amplicons. Precise repair at A but not B, was shown by properly sized PCR A amplicon and lack of or incorrect size of PCR B amplicon. Precise repair at B but not A was indicated by a properly sized PCR B amplicon and lack of or incorrect size of PCR A amplicon. Analysis by Sanger sequencing of
homozygous strains with each class of repair from the three PAM orientations confirmed that the properly sized PCR amplicons reflected precise repair at a DSB junction.

### Examining the effects of mating on genome editing efficiency

Matings were set up between males and hermaphrodites with genotypes shown in Figures 10-12, and S5 by placing two L4 hermaphrodites and 10 males on NG agar plates with small (~0.5 cm diameter) spots of OP50 bacteria. Worms were allowed to mate at 20° for 24 hr prior to microinjection. Young adult hermaphrodites were then transferred to new NG agar plates without bacteria prior to mounting for microinjection. Swollen spermathecae visible at the time of injection indicated that hermaphrodites had mated successfully. After microinjection and recovery at 20° for 3 hr, P0 worms were transferred to individual NG agar/OP50 plates and incubated for three days at 25°. Presence of F1 male offspring provided further evidence that mating had occurred. Both Rol and/or Dpy hermaphrodites and males produced by successfully mated mothers were transferred to individual wells of a 96-well plate for lysis and genotyping, as described above.

In our experience using *dpy*-*10* as a co-conversion marker, half or more of injected self-fertile P0 hermaphrodites yielded a high frequency of F1 progeny with Dpy or Rol phenotypes. However, for experiments in Figures 11 and 12 in which hermaphrodites were mated with males prior to injection and the *dpy*-*10* allele from only one parent could be targeted by Cas9, only a quarter or less of the injected P0 animals produced F1 progeny with Dpy or Rol phenotypes.

The genomic feature that permitted selective targeting of maternal vs. paternal chromosomes was a single base pair change in a critical region of the spacer DNA required for pairing with the guide RNA. When targeting X-specific loci using a guide that could only pair with the male sperm-specific allele, F1 males produced by the cross were examined by DNA sequence analysis to determine whether the inherited maternal X chromosome had been edited via the mismatched guide. We found that male progeny lacked edited X chromosomes, indicating that the single-base mismatch in the spacer seed sequence of the maternal genome prevented its cleavage.

### Strategy for scoring inter-homology HDR events

For Figures 11 and 12, classifying DNA changes as the result of inter-homolog HDR events required distinguishing cross-progeny from self-progeny. By experimental design, animals that arose from interhomolog HDR events would be Rol or Dpy hermaphrodites that carried homozygous *sex*-*1* GG alleles. However, in some crosses, Rol or Dpy *sex*-*1* GG homozygous animals could have been self-progeny of *sex*-*1* GG homozygous hermaphrodites that failed to mate with *sex*-*1* AG males. Thus, we only considered Rol or Dpy *sex*-*1* GG homozygous animals to be the result of inter-homolog HDR events if they were cross-progeny.

For Figure 11A, 11D and 12A, the RNA guides permitted only the *dpy*-*10* AG allele contributed from the male sperm to be edited. Therefore, all Rol or Dpy animals had to be cross-progeny, and Rol or Dpy *sex*-*1* GG homozygous animals should have arisen from cleavage of the *sex*-*1* AG allele and repair using the *sex*-*1* GG allele contributed by the other parent.

In Figure 11B and 12B, the editable *dpy*-*10* AG allele was contributed by the mother. Thus, all Rol or Dpy hermaphrodites that were homozygous for the paternal *sex*-*1* GG allele could be classified as cross-progeny that underwent inter-homolog HDR events.

In Figure 11C, the frequency of inter-homolog HDR events was estimated by assuming that all assayed F1 Dpy or Rol progeny were cross-progeny. We considered this assumption to be valid based on our results from experiments in panels A, B and D, in which cross-progeny could be identified unambiguously. In those experiments, all assayed Dpy or Rol progeny were cross-progeny, suggesting that all matings in the experiments of Figure 11 were fully successful. Several guidelines helped ensure that matings were successful, and the assayed Dpy or Rol progeny were cross-progeny. First, when mounting P0 worms for microinjection, the mated hermaphrodites were examined using differential interference contrast microscopy to confirm successful matings by the presence of engorged spermathecas. Second, prior to assaying Rol or Dpy F1 animals, the F1 offspring were examined to confirm that nearly half of the F1 offspring were males, an indication of fully successful mating prior to microinjection.

For Figure 11, A and B and Figure 12, A and B, repair events classified as inter-homolog HDR required that the cleaved target in the haploid genome of one gamete be repaired using the homologous chromosome in the haploid genome of the other gamete, thereby generating an embryo that was homozygous for a SNP in *sex*-*1*. Primers that anneal ~200 bases from the *sex*-*1* SNP were used for PCR and Sanger sequencing to confirm the genotype of the offspring. It was formally possible that a cleaved chromosome could have been repaired imprecisely to yield a deletion that would have prevented the PCR primers from annealing to the deleted sequences in the targeted *sex*-*1* locus. In that case, only sequences from the uncut chromosome would be amplified, and the *sex*-*1* locus would be incorrectly classified as homozygous for the SNP due to inter-homolog HDR. To rule out such false positive results, 10 F1 progeny that appeared to have undergone inter-homolog HDR at *sex*-*1* were allowed to produce F2 progeny. Sixteen F2 progeny from each of the 10 different F1 animals were lysed and their DNA sequenced to confirm that the edits were indeed homozygous. We found that PCR never failed to amplify the loci, and all the DNA sequences were consistent with inter-homolog HDR events. In these *sex*-*1* experiments, deletions in the vicinity of the SNP do not cause any lethality, and we found no death among F2 progeny. Thus, our analysis of F2 progeny would not have missed false-positive inter-homolog HDR events due to lethality.

## Results and Discussion

### Imprecise repair at Cas9 cleavage sites is asymmetric, favoring changes 5′ of the PAM

Analysis of imprecise repair outcomes at Cas9 DSBs in *sex*-*1* and *lir*-*2* revealed a polarity to the insertion and deletion of sequences that occurred in the absence of exogenous repair templates. Changes were preferentially introduced 5′ of the PAM (Figure 1). Even when imprecise repair caused deletions 3′ of the PAM, the deletions were substantially shorter in the 3′ direction than in the 5′ direction on the same DNA strand. The propensity of repair to favor changes 5′ of the PAM that is true for nematodes has been found in mammals as well (Mali *et al*. 2013b; Van Overbeek *et al*. 2016; Richardson *et al*. 2018), providing the opportunity to use *C. elegans* as a model to understand general rules underlying such directionality and apply them to achieve changes in desired locations.

### Precise repair from homologous single-stranded oligonucleotide templates is directional, and its efficiency is dictated by the choice of repair template strand

To determine whether homology-directed repair is also asymmetric, we compared repair outcomes using single-stranded oligonucleotide templates that matched either the protospacer strand or the spacer strand of three different loci (*lir*-*2*, *sex*-*1*, and Chr. I site) (Figure 2, A and B). For both classes of repair templates, two adjacent single nucleotide polymorphisms (SNPs) were positioned at a site corresponding to the chromosomal DSB site. These two adjacent SNPs constitute polymorphism b. Additional SNPs were also positioned 10 nt from the DSB in both 5′ and 3′ directions (Table S2). As a convention, we defined the positions of edits at endogenous loci relative to the PAM on the protospacer strand.

For both protospacer and spacer strand repair templates, insertion of polymorphisms at DSB sites was efficient at all three endogenous loci [15%-47% of *dpy*-*10* mutants], but a striking preference was found for insertion of SNPs 5′ of the PAM when the protospacer strand was used for the repair template. As shown in Figure 2A, 12%-37% of the *dpy*-*10* mutants had a SNP inserted 5′ of the PAM from the protospacer template but 0% of worms had a SNP inserted 3′ of the PAM. However, if the spacer strand was used for the repair template, insertion of a SNP 3′ of the PAM was strongly favored. We found that 6%-13% of *dpy*-*10* mutants had a 3′ polymorphism inserted from the spacer strand template, but only 0%-4% of *dpy*-*10* mutants had a 5′ polymorphism inserted (Figure 2B). Because we detected both imprecise repair and homology-directed repair at high frequency at the predicted Cas9 cleavage sites, we are certain that the lack of homology-directed repair in either the 5′ or 3′ direction was not caused by the failure of Cas9 to cleave DNA (Figure 2, A and B).

The dramatic polarity in repair can be explained by a repair process that proceeds via a synthesis-dependent strand annealing mechanism (SDSA), as diagrammed in the models of Figure 2C (Sekelsky 2017). The 3′ end of the cleaved spacer strand in the endogenous locus can anneal readily with the complementary protospacer strand repair template and prime DNA synthesis from the template, thereby incorporating polymorphisms only at the DSB and 5′ of the PAM (Figure 2C left). Similarly, the 3′ end of the cleaved protospacer strand in the endogenous locus can anneal with the complementary spacer strand repair template and prime DNA synthesis from the template, thereby incorporating polymorphisms only at the DSB and 3′ of the PAM (Figure 2C right). This SDSA model is rigorously supported by the repair outcomes from our two complementary sets of experiments and has been proposed previously by others in the context of repair from a single-stranded break generated by a Cas9 mutant variant having only one of its two single-stranded nickases (Davis and Maizels 2014; Davis and Maizels 2016) and in the context of repair from Cas9 DSBs (Kan *et al*. 2017; Paix *et al*. 2017)) and meganucleases (Kan *et al*. 2014).

The occasional insertion of SNPs 5′ of the PAM when a spacer strand repair template was used can be explained by the 3′ exonucleolytic activity for the Cas9 nickase RuvC (see model in Figure S1) (Jinek *et al*. 2012; Jiang *et al*. 2015; Zuo and Liu 2016; Stephenson *et al*. 2018). This nickase resects single-stranded DNA *in vitro* up to 10 nucleotides in the 3′ - 5′ direction on the strand it nicks (Stephenson *et al*. 2018). Since polymorphism a is only 10 bp from the DSB, the RuvC nickase should occasionally resect the endogenous DNA just past the location of the polymorphism and permit primer extension from the spacer repair template to incorporate this polymorphism into the endogenous locus at low frequency, as we observed.

We addressed two additional issues about DSB repair from single-stranded repair templates: (1) whether the presence of polymorphisms on both sides of the DSB, as in Figure 2, blocks the template from being used efficiently for repair and (2) whether all polymorphisms within 30 bp of a DSB will be inserted with similar frequency relative to those closest to the DSB. If repair is very local, the repair process may not insert the more distant changes in the repair template.

We addressed both topics by using single-stranded repair templates designed to insert polymorphisms immediately adjacent to the chromosomal DSB site (polymorphism d) and also in one direction from the DSB, in 10 nt intervals (Figure 3). Similar to the pattern observed using repair templates with symmetric changes, repair templates corresponding to the protospacer strand resulted in efficient insertion of polymorphisms directly over the DSB (30% and 26%) and 5′ of the PAM (14%), but not 3′ of the PAM (0%) (Figure 3A). Remarkably, incorporation of all four changes in the 5′ direction was more efficient (12%) than incorporation of only a subset of the changes (2%) (*P* < 0.02, chi-square). In contrast, repair templates corresponding to the spacer stand (Figure 3B) resulted in efficient insertion of polymorphisms at the DSB (39% and 16%) and 3′ of the PAM (22%), but not 5’ (0%). Insertion of all four changes in the 3′ direction (11%) was on par with insertion of only a subset of changes (11%), in contrast to the protospacer template result.

Several generalizations emerged from this set of experiments. Insertion of polymorphisms specifically at the DSB site is highly efficient regardless of whether the single-stranded repair template corresponds to the protospacer strand or spacer strand. However, insertion of SNPs 5′ of the PAM is far more efficient if the repair template corresponds to the protospacer stand, while insertion of SNPs 3′ of the PAM is much more efficient if the repair template corresponds to the spacer strand. Therefore, the choice of repair template strand is critical for ensuring the insertion of polymorphisms in the desired locations. Finally, when SNPs were inserted either 5’ or 3’ of the PAM, a tendency prevailed to incorporate all of the SNPs in one direction, not just a subset of the SNPs, particularly for the protospacer repair template.

The precise repair outcomes we observed were independent of whether the strand of the repair template was non-coding (*sex*-*1*, *lir*-*2*), coding (*zen*-*4*, Table S2, Materials and Methods), or intergenic (site on chromosome I). Others have suggested that successful insertion of polymorphisms was higher if the coding strand of the gene was used as the repair template rather than the non-coding strand (Katic *et al*. 2015). However, our re-examination of the published repair outcomes revealed that the polarity can be explained by the location of the polymorphism relative to the PAM and is therefore consistent with the strong trend in our experiments.

### For homology-directed repair at a single Cas9 DSB, a double-stranded repair template was less efficient than a single-stranded template

Optimization of genome-editing strategies requires a comparative analysis of HDR efficiency with single-stranded vs. double-stranded repair templates. We initiated this comparison by repeating the experiment of Figure 2A with a linear double-stranded repair template that was identical in sequence to the 100 nt *sex*-*1* single-stranded HDR template. To our surprise, we observed no insertion of SNPs at the *sex*-*1* locus in 144 animals that were edited at the *dpy*-*10* locus (Figure 4A). Instead, 75% of the *dpy*-*10* mutants had undergone imprecise repair at the *sex*-*1* locus, indicating that Cas9 cleavage was effective but that HDR from the double-stranded template was completely ineffective. Furthermore, the lack of HDR was not caused by the double-stranded repair template being destroyed by Cas9 cleavage, because the template carried SNPs in the sequence required by the guide RNA (Materials and Methods and subsequent experiments).

In a second experiment, we used a 1 kb linear double-stranded repair template to *sex*-*1* that included the same SNPs as the 100 bp template but replaced the 40 bp homology arms with 490 bp arms (Figure 4A). Results with this template were more successful but still not robust. The frequency of imprecise repair was reduced to 39%, and the frequency of HDR elevated: 4% of *dpy*-*10* mutants had SNP insertions at the DSB site, and 1-2% also had SNP insertions on either side of the DSB, but not both sides. Thus, repair with linear double-stranded repair templates was considerably less efficient than repair with single-stranded templates.

A third experiment assessed the maximum distance from the DSB that SNPs were inserted into *lir*-*2* from a double-stranded repair template (Figure 4B). The 4 kb HDR template was embedded in a circular plasmid, had 500 bp homology arms, a SNP that disrupted the PAM (hence Cas9 binding) and 10 polymorphisms positioned at ~100 bp intervals on both sides of the DSB. Each polymorphism had 1-4 bp substitutions that created a *Hind*III site for monitoring HDR. HDR repair outcomes were infrequent (6% of 380 *dpy*-*10* mutants) and exhibited a striking polarity in which insertions occurred up to 900 bp from the DSB but primarily in only one direction, 5′ of the PAM. The exception was a polymorphism that was inserted 100 bp from the DSB in the opposite direction. Thus, the strategy of using a single Cas9 cleavage site and a double-stranded repair template to edit the genome yielded only a low frequency of HDR events. These events were too directional and limited to permit insertion of large DNA fragments with multiple polymorphisms.

A final set of experiments assessed whether a large non-homologous DNA fragment (9300 bp) could be inserted into the genome at a single DSB site. We evaluated double-stranded repair templates with three different sets of 500 bp homology arms. We found that 0% of 576 edited animals had an insert of the 9300 bp DNA fragment, thus reinforcing the conclusion that large DNA fragments are difficult to insert into the genome using only one DSB (Figures S2 and 5A).

### Efficient strategies to insert large DNA fragments at wide-ranging distances from DSBs

To overcome the strong polarity in HDR events repaired from double-stranded templates, the inefficiency in inserting polymorphisms that are distant from a DSB, and the inefficiency in inserting large non-homologous fragments into the genome, we explored whether two nearby DSBs made using two different guide RNAs simultaneously would facilitate insertion of long DNA fragments (Figure 5A). Two DSBs have been used to make chromosomal rearrangements, including inversions translocations, and large deletions (Blasco *et al*. 2014; Chen *et al*. 2014; Choi and Meyerson 2014; Ghezraoui *et al*. 2014; Maddalo *et al*. 2014; Paix *et al*. 2014; Iwata *et al*. 2016; Dejima *et al*. 2018). We found that adding a second DSB at a distance of 340 bp from the initial DSB (Figures 5A and S2) enabled us to insert a 9300 bp fragment from a double stranded template with reasonable efficiency (5%) (Figures 5B).

This double DSB strategy also worked well using a variety of linear and circular double-stranded repair templates with 500 bp homology arms to insert fluorescent reporters: 21% *dpy*-*10* mutants had precise insertions of 8221 bp, 17% had precise insertions of 5060 bp, and 36% had precise insertions of 2138 bp (Figure 5B). In all cases, the DNA between DSBs (340 bp or 98 bp) had been deleted, and the desired DNA inserted. Our approach using two nearby DSBs to incorporate long DNAs, is sufficiently robust and precise that selection strategies are not needed to enrich the number of candidates to screen for full-length insertions.

A slight variation on this strategy enabled the insertion of desired DNA at long distances from DSBs without deleting any intervening DNA. The variation is highly useful when the target sites for editing lack nearby sequences needed for effective Cas9 guides or when a series of different substitutions are desired simultaneously along the length of DNA (Figure 5C,D). The adaptation depends solely on the design of the repair template. For example, we inserted a 1275 bp fragment with good efficiency in various locations within the 2826 bp region of DNA between two DSBs in *xol*-*1* simply by using different 4101 kb HDR templates that included the entire 2826 bp homologous region with the 1275 bp non-homologous insert placed at various locations within the homologous region (Figure 5C). The non-homologous DNA could be inserted as far as 1968 bp from one of the DSBs. The purpose was to insert 24 copies of the binding site for the MS2 phage coat protein into different introns for the goal of performing live imaging of transcription.

By a similar strategy, we deleted three short regions (25 bp-37 bp) within an intron of *xol*-*1* and made three bp substitutions at 5 different locations within the intron to define the binding sites used by an RNA binding protein *in vivo*. In the latter experiment, all the desired SNPs were incorporated into each of the edited introns. Thus, the high efficiency of inserting long, full-length stretches of DNA into the genome using two DSBs provides the flexibility to perform genome editing at multiple sites in a large region of genomic DNA.

Others reported that repair template homology arms greater than 50 bp can reduce the efficiency of HDR (Paix *et al*. 2014). We have obtained good HDR efficiency with 500 bp homology arms but also tested 50 bp homology arms to ask whether the efficiency could be improved. We found it was not (Figure 5D).

### For HDR induced by two Cas9 cleavage sites, relative orientation of PAMs influences efficiency of HDR

Since successful genome editing using a single Cas9 cleavage site required a specific PAM orientation relative to the site of editing and the repair template strand, we explored the significance of PAM orientation on HDR efficiency when two simultaneous DSBs were employed. Cas9 binds PAMs tightly both *in vitro* and *in vivo* and remains bound after DNA cleavage. Each DSB generates two DNA ends: the end with the PAM and the end without the PAM. These two unique DNA ends are likely to be processed differently during DSB repair and may support different rates of homology directed repair.

To understand the effect of PAM orientation on HDR using two DSBs, we examined repair rates at the same genomic locus in three strains that differed only by the orientation of PAMs at the two Cas9 target sequences. The PAMs were arranged in an OUT/OUT, IN/IN, or IN/OUT orientation (Figure 6). The PAM OUT orientation left the PAM on the chromosome end after DNA excision. The PAM IN orientation placed the PAM within the DNA sequence excised from the chromosome. Since the three strains harbor the same two unique Cas9 target sequences, strain-specific differences in repair frequencies should reflect differences in PAM orientations and not differences in Cas9 recognition and cleavage efficiency.

Orienting both PAMs outward (OUT/OUT) resulted in the most efficient rate of homology directed repair: 21% of *dpy*-*10* hermaphrodites had the templated changes (Figure 6). In this OUT/OUT orientation, the PAMs would be retained on the chromosome ends after DNA cleavage. In the other extreme orientation, the IN/IN orientation, 12% of the *dpy*-*10* mutants had the templated sequences. For this orientation, the PAMs would be on the excised piece of DNA. The least efficient repair occurred with one PAM in the IN orientation and the other in the OUT orientation (IN/OUT), resulting in one PAM on the excised DNA end and the other on the chromosome end. Only 4% of Rol or Dpy mutants had the precise insert. A similar HDR pattern held true for the much larger repair template used in Figure S4 in which HDR from the IN/IN orientation was 19% but only 7% from the IN/OUT orientation.

No permutation of PAM orientations resulted in the total failure of HDR for the fully matched set of experiments in Figure 6, or the more random experiments in Figure 5. Thus, although the most successful strategy for efficient HDR insertions is to use Cas9 targets that position both PAMs in the OUT orientation, other orientations can also work, just not as effectively. The low efficiency of achieving insertions with either the IN/OUT or OUT/IN orientation of PAMs likely accounts in part for the difficulty in inserting large segments of DNA using a single Cas9 guide and hence only one DSB, which naturally causes the cleaved ends to have an IN/OUT orientation.

A relevant set of experiments involving different PAM orientations explored the effect of overhangs made from Cas9 variants, each with one nickase activity, either RuvC or HNH (Bothmer *et al*. 2017). Consistent with our results, the experiments showed that loci with two overhangs having the PAM OUT/OUT orientation gave a higher frequency of imprecise repair than loci with two overhangs having the PAM IN/IN orientation.

Our experiments recovered not only animals with precisely repaired DNA at both DSBs, but also animals with precisely repaired DNA at one DSB and imprecisely repaired DNA at the other DSB (Figure 6). No pattern emerged that correlated the IN or OUT PAM orientation with precise versus imprecise repair at either of the two DSB junctions.

### Optimizing guide RNA design and Cas9 delivery

Efficient genome editing requires optimization of the guide RNA design to ensure a high frequency of Cas9-induced DSBs. We showed previously for *C. elegans* that guide RNAs having a GG sequence at the 3′ end of their target-specific sequences were the most effective for achieving targeted mutagenesis (Farboud and Meyer 2015). We re-visited the importance of the GG guide sequence, because we changed our protocol for delivering Cas9 to germline nuclei from injecting DNA expression vectors encoding Cas9 and guide RNAs to the more effective protocol of injecting pre-assembled Cas9/guide RNA ribonucleotide particles (RNPs). Using both delivery protocols, we compared directly the efficiency of genome editing using guides having GG, AG, CG, or UG at their 3′ ends (Figure 7). To do so, we first performed genome editing to make four isogenic strains in which *sex*-*1* genomic DNA had been changed by a single nucleotide substitution to convert the penultimate 3′ G of the protospacer to either A, C, or T.

With the DNA expression vector delivery approach, only the *sex*-*1* GG RNA guide induced imprecise repair (68%) at the endogenous *sex*-*1* locus of animals edited at *dpy*-*10*. In contrast, RNP injections were more permissive, but the GG RNA guide was still far more effective at genome editing: while 71% of GG RNA guides induced imprecise repair at the *sex*-*1* locus, only 28% of AG guides, 18% of CG guides, and 12% of UG guides induced *sex*-*1* changes. Thus, while genome editing in *C. elegans*, as in other organisms, can be accomplished with guides lacking a G as the penultimate 3′ nucleotide of the target-specific sequence, the efficiency of mutagenesis in *C. elegans* is much higher if the guides have a penultimate G.

### Efficient co-conversion marker to edit genomes of evolutionarily diverged nematodes

Most co-conversion markers for *C. elegans* (*dpy*-*10*, *rol*-*6*, *sqt*-*1*) reside on chromosome II, making it inconvenient to edit another gene on chromosome II because the edited gene will have an unwanted linked visible marker (Arribere *et al*. 2014). Furthermore, mutations in neither *rol*-*6* nor *dpy*-*10* have strong dominant phenotypes in *C. briggsae*, thus limiting their utility as co-conversion markers in this diverged sister species (15-30 MYR).

To overcome both problems, we developed *ben*-*1* as a co-conversion marker effective for both *C. elegans* and *C. briggsae* (Figure 8, A and B). *ben*-*1*, a gene on chromosome III, encodes β-tubulin. While developing *C. elegans* genome editing protocols, we showed previously that *ben*-*1* loss-of-function mutations made using zinc-finger nucleases and TALENS were dominant for conferring resistance to the drug benomyl (Wood *et al*. 2011; Lo *et al*. 2013). Benomyl binds to β-tubulin, inhibits microtubule polymerization, and causes wild-type animals to be slow growing and paralyzed. *ben*-*1* loss-of-function mutations enable *C. elegans* to be mobile in the presence of benomyl (Driscoll *et al*. 1989).

**Figure 8.**
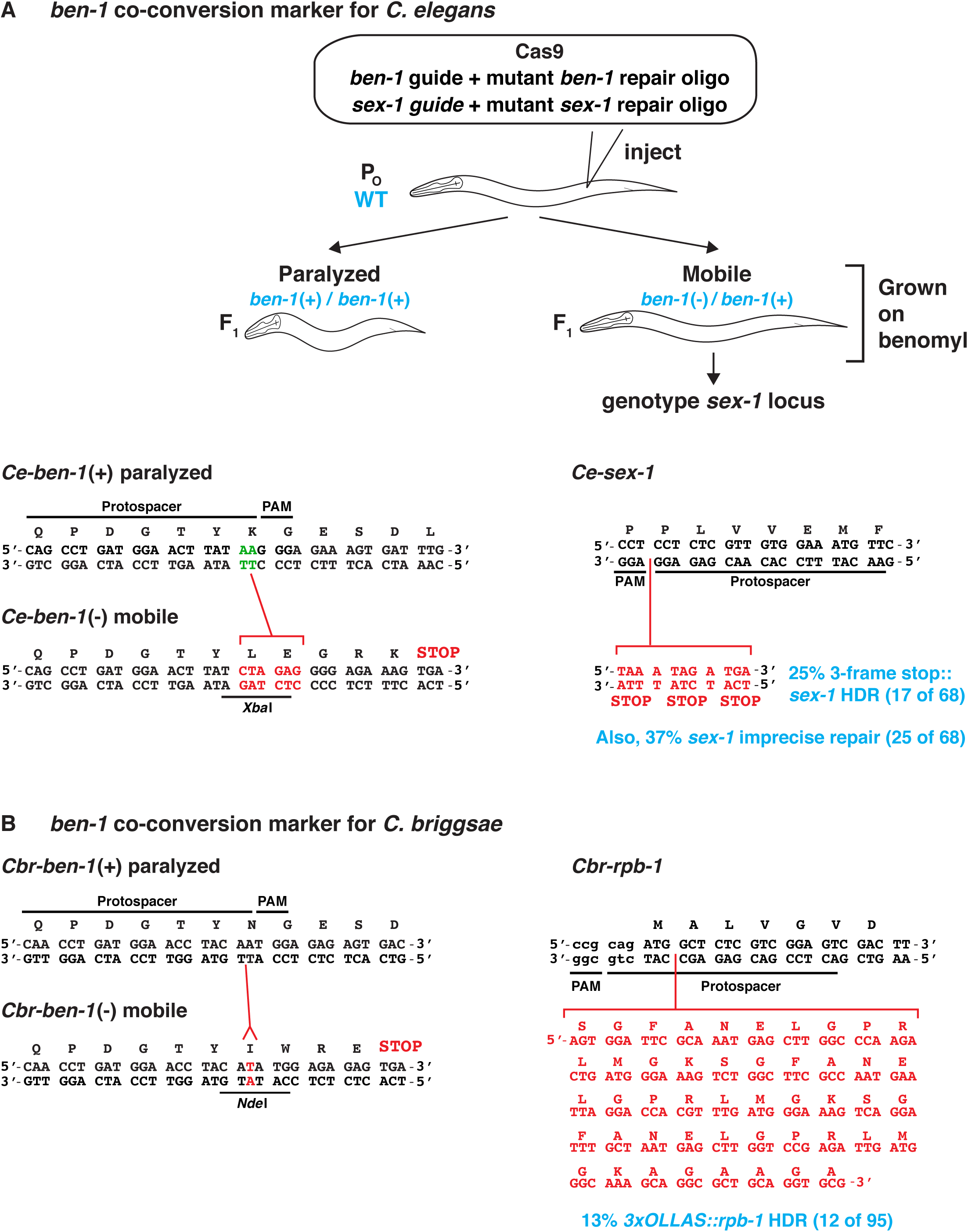
Efficient co-conversion marker to edit genomes of diverged nematode species Mutations in the highly conserved gene *ben*-*1*, which encodes β-tubulin, confer resistance to benomyl, a drug that binds to β-tubulin, inhibits microtubule polymerization, and causes genetically wild-type animals to be slow growing and paralyzed. Loss-of-function mutations in *ben*-*1* confer dominant benomyl resistance in both *C. elegans* and *C. briggsae*. A) In *C. elegans*, the repair template created an *Xba*I restriction site adjacent to the PAM by converting an AA to CTAGAG, resulting in an in-frame stop codon that caused premature translation termination. (B) In *C. briggsae*, the repair template created an *Nde*I site by inserting a T adjacent to the PAM, causing an in-frame translation termination stop codon. In both species, the repair template altered protospacer sequences adjacent to the PAM, preventing the *ben*-*1* mutants from being cleaved by Cas9. Successful editing of *C. elegans sex*-*1* and *C. briggsae rpb*-*1* established *ben*-*1* as an effective co-conversion marker.

The *ben*-*1* co-conversion strategy was highly efficient in *C. elegans* experiments (Figure 8A). To develop *ben*-*1* as a co-conversion marker we used an AA guide for *ben*-*1* and a single-stranded oligonucleotide HDR repair template to convert an AA to CTAGAG, thereby creating a new *Xba1* site and a premature in-frame translation stop codon downstream (Figure 8A). For measuring co-conversion, we targeted *sex*-*1* with a GG guide and a single-stranded oligonucleotide repair template to insert 11 bp of new sequences, thereby creating stop codons in all three reading frames to prevent translation of the SEX-1 activation function 2 domain (AF2). We found that 62% of mobile *ben*-*1* mutant animals had either an imprecise (37%) or precise (25%) repair event at the *sex*-*1* locus in co-conversion experiments (Figure 8A)

The *ben*-*1* co-conversation strategy was also efficient in *C. briggsae* experiments. We mutated *C. briggsae ben*-*1* using an AA guide and a single-stranded oligonucleotide HDR repair template to insert a T adjacent to the PAM, thereby creating a new *NdeI* site and a premature in-frame translation stop codon downstream. As in *C. elegans*, the resulting loss-of-function *ben*-*1* mutations were dominant in *C. briggsae*, enabling hermaphrodites to be mobile in the presence of benomyl (Figure 8A and Materials and Methods). In co-conversion experiments, we found that 13% of mobile benomyl-resistant animals had an HDR repair event that precisely introduced a 144 bp insert to create a *3xOLLAS* tag in the *rpb*-*1* locus. This result established *ben*-*1* as a good co-conversion marker for *C. briggsae*.

### Selectable co-conversion marker for C. elegans

To achieve greater ease and efficiency in identifying candidates with genome editing events, we engineered the gene *zen*-*4* to generate a co-conversion marker that would be strongly selectable by permitting reversion of a temperature-sensitive lethal mutant phenotype to wild-type. Amino acid substitution D520N in ZEN-4 causes rapid, temperature-sensitive embryonic lethality within 30 minutes after *zen*-*4* mutants are shifted from the 15° permissive temperature to the 25° non-permissive temperature (Jantsch-Plunger *et al*. 2000; Severson *et al*. 2000). *zen*-*4* (chromosome IV) encodes a member of the kinesin-6 superfamily of plus-end-directed microtubule motors that is required for polar body extrusion during meiotic divisions, for formation and maintenance of spindle midzone microtubules, and for completion of cytokinesis after mitosis (Raich *et al*. 1998; Jantsch-Plunger *et al*. 2000; Severson *et al*. 2000). We used genome editing to create a temperature-sensitive lethal nematode strain carrying this temperature-sensitive lethal *zen*-*4* allele optimized for use as a co-conversion marker.

The lethal mutant strain *zen*-*4(cle10ts)* carried three changes (Figure 9 and Figure S3). The first change was a GAC to AAC transition at codon 520 to create the temperature-sensitive lethal allele. The second was a CGA to CGG synonomous transition at codon 523 to create a PAM without any change in protein sequence. The third was an *Alu1* restriction site introduced by converting codon 519 from GCA to GCT, again with no animo acid change. The single-stranded oligonucleotide repair template would revert *zen*-*4(cle10*ts) to the wild-type allele, permitting successfully edited animals to be viable at the non-permissive temperature and hence analyzed for co-editing of the primary target gene. Using this strategy, the *Alu*I site would also be eliminated, enabling viable *zen*-*4*(+) edited animals to be distinguished from true revertants by a simple PCR assay. The PAM created by the lethal *cle10*ts mutation would also be lost, thereby preventing the repaired *zen*-*4* allele from being cleaved again by Cas9 and damaged. This selection scheme has an advantage over the selectable co-conversion *pha*-*1* scheme in not leaving a SNP behind in the genome (Ward 2015).

**Figure 9.**
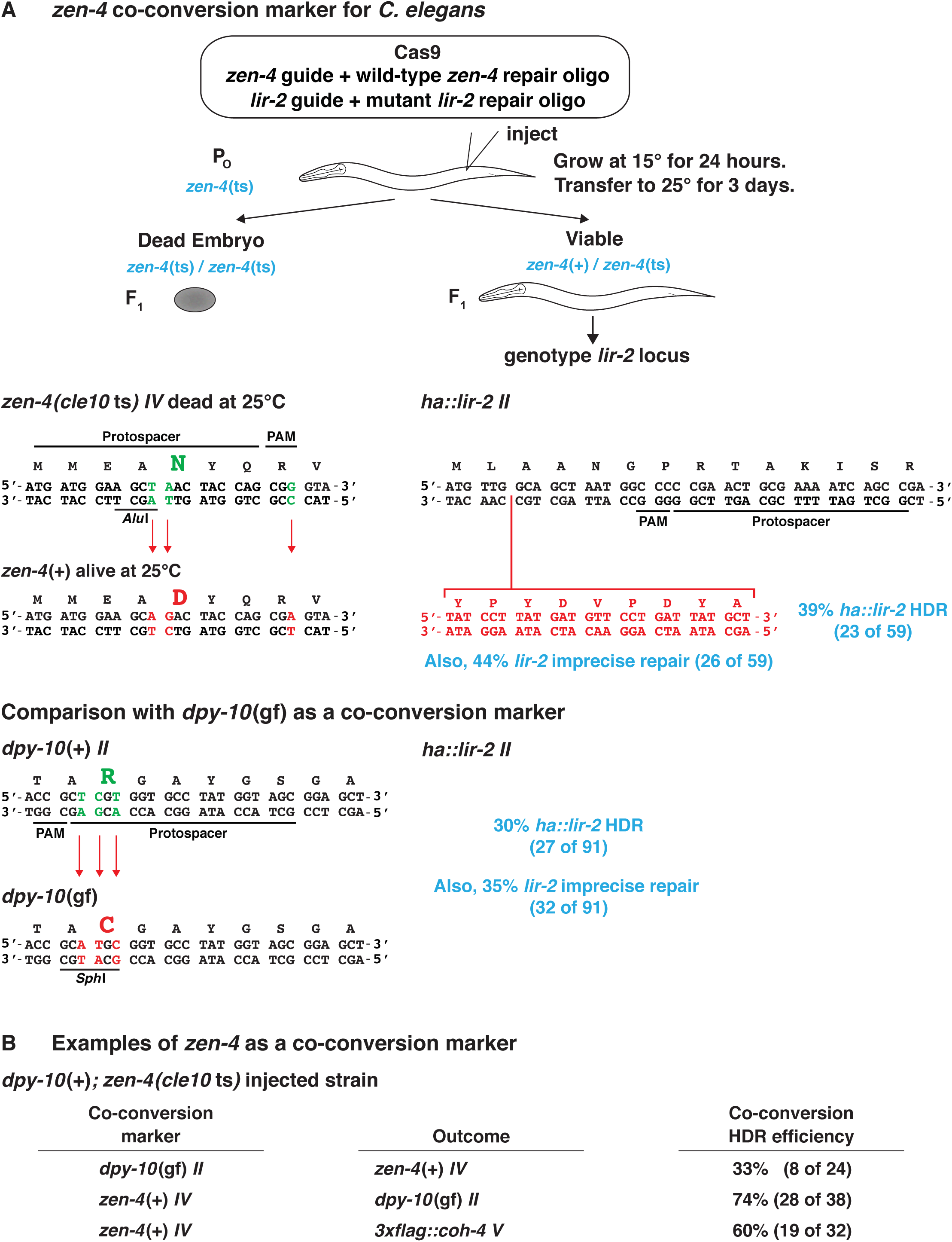
Selectable co-conversion marker for *C. elegans* (A) *zen*-*4*(+) produces an essential plus-end-directed microtubule motor and acts as an efficient co-conversion marker to revert the rapid, temperature-sensitive lethality caused by a defective ZEN-4 protein with amino acid substitution D520N produced by *zen*-*4*(*cle10*ts) mutants. Viable *zen*-*4*(+) progeny are scored for editing in genes of choice. (B) Successful editing of *lir*-*2*, *dpy*-*10*, and *coh*-*4* established *zen*-*4* as an effective co-conversion marker. Comparison of editing efficiency using *dpy*-*10* and *zen*-*4* as co-conversion markers is also presented.

We tested the efficiency of *zen*-*4* as a co-conversion marker by comparing the frequency of inserting DNA encoding a hemagglutinin (HA) tag into the *lir*-*2* gene using *zen*-*4* or *dpy*-*10* as a co-conversion marker. Of viable *zen*-*4*(+) animals, 39% had the *ha::lir*-*2* insertion, and 44% had imprecise repair at *lir*-*2* (Figure 9A). The 83% overall success rate at *lir*-*2* was more favorable than the 65% rate with *dpy*-*10* as the co-conversion marker (30% with the *ha::lir*-*2* insertion and 35% with imprecise repair). This result suggested that *zen*-*4* editing is less efficient than *dpy*-*10* editing, causing the relative frequency of edits at desired loci to be higher in *zen*-*4*(+) animals than in *dpy*-*10* mutants. This hypothesis was validated by our result that only 33% of *dpy*-*10* mutants were converted to *zen*-*4*(+) (Figure 9B). Decreased editing efficiency at *zen*-*4* is likely due to the need for precise HDR repair to yield a wild-type *zen*-*4*(+) phenotype, but both precise and imprecise repair at *dpy*-*10* yield obvious phenotypes. The decreased editing efficiency and the strong selection for viability make *zen*-*4* a valuable co-conversion marker.

*zen*-*4* was highly successful as a co-conversion marker for editing other loci (Figure 9B). For example, 60% of *zen*-*4*(+) edited animals acquired a DNA insertion encoding a FLAG tag in the cohesin subunit gene *coh*-*4*. Also, 74% of *zen*-*4*(+) edited animals acquired a *dpy*-*10* mutation.

### Efficient genome editing occurs in embryos

Thus far, we have evaluated factors that influence the success of genome editing in self-fertile hermaphrodites. In these hermaphrodites, Cas9 RNPs and repair templates were introduced into the gonad syncytium of young adults that had already produced a full complement of sperm but still had hundreds of prophase nuclei destined to become cellularized and mature into oocytes. For fertilization to occur, the Cas9-exposed oocytes had to pass through the spermatheca, the organ that stores mature ameboid spermatozoa derived from either self-fertile hermaphrodites or males during male/hermaphrodite mating. In principle, Cas9-induced cleavage and repair events could occur at any time during meiotic prophase or later during embryogenesis, either before or after oocyte and sperm pronuclei had formed and fused. To distinguish between these possibilities, we launched a series of experiments to evaluate (i) the frequency and type (precise vs. imprecise) of DNA editing in embryos versus that in meiotic prophase; (ii) the efficiency of editing genomes from male sperm versus genomes from hermaphrodite sperm or oocytes; and (iii) the effect of copulation *per se on* genome editing.

We first addressed whether differences in editing efficiency were detectable between progeny from self-fertile hermaphrodites versus progeny from hermaphrodites mated for 24 hours by wild-type males (Figure 10, A and B). Using *dpy*-*10* as a co-conversion marker, we found that the frequency of indels in the *aft*-*2* II gene of *dpy*-*10* mutants rose to almost twice the level in the outcrossed progeny of mated hermaphrodites (61%) versus the self-progeny of unmated hermaphrodites (35%) (*P* = 0.001, chi-square).

**Figure 10.**
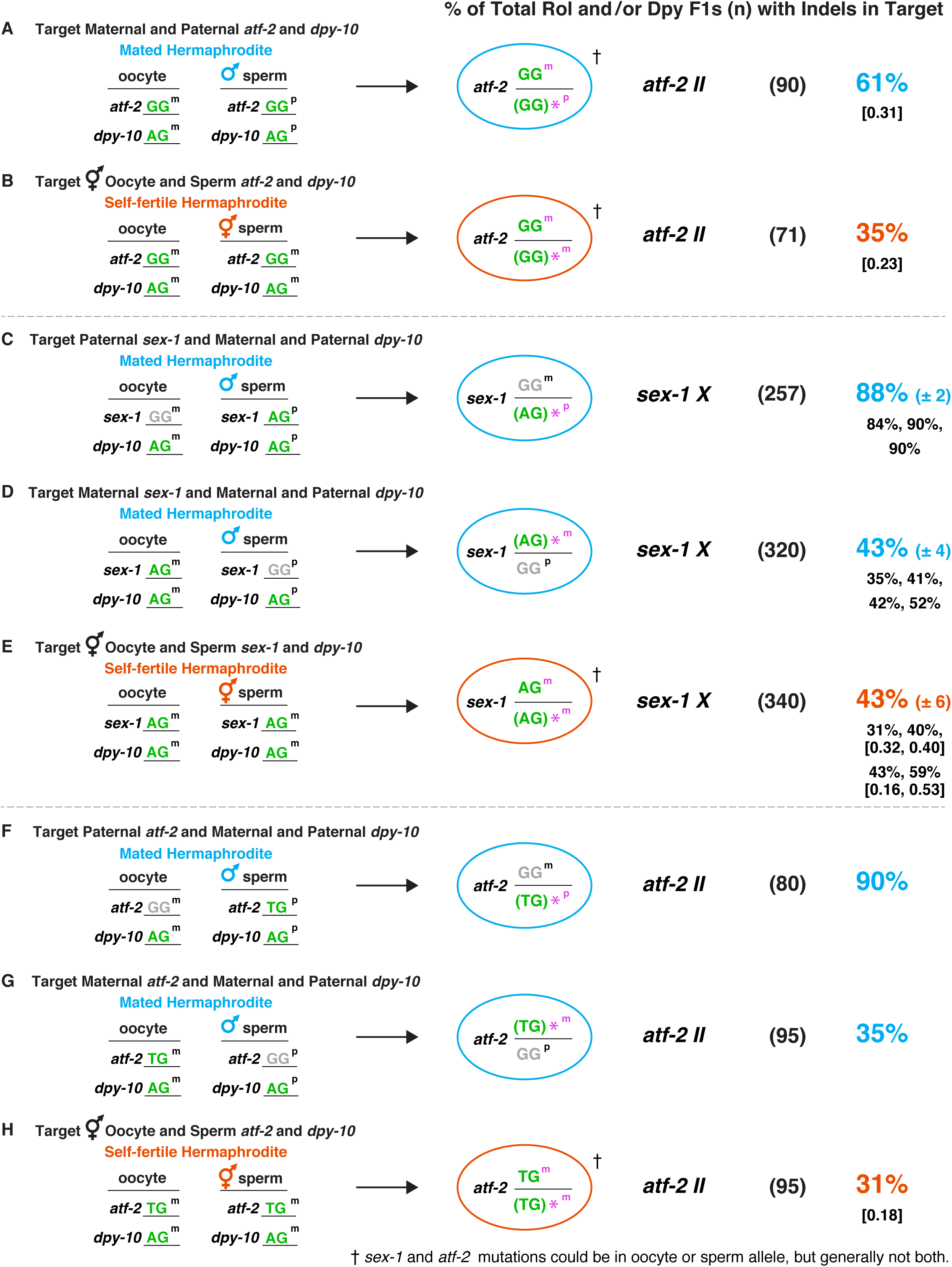
Efficient genome editing occurs in embryos of mated hermaphrodites (A and B) Quantification of indels at *aft*-*2 II* in outcrossed progeny of mated hermaphrodites (A) versus self-progeny of self-fertile hermaphrodites (B) shows a higher indel frequency in outcrossed progeny (*P* </italic>= 0.001, chi-square). (C-H) Comparison is shown for the frequency of indels at *sex*-*1* X (C-E) and *atf-2* II (F-H) in genomes derived from (i) male sperm (paternal genome) and (ii) hermaphrodite oocytes (maternal genome) in outcrossed progeny from mated hermaphrodites and from (iii) sperm and/or oocytes in self-fertile hermaphrodites. For *sex*-*1* and *atf*-*2*, the frequency of indels in the paternal allele of progeny from mated hermaphrodites was at least twice that for the identical maternal allele (*P* < 10^−5^, chi-square) in progeny of mated hermaphrodites and for the identical alleles in progeny of self-fertile hermaphrodites (*P* < 10^−5^, chi-square), even though the male sperm genome had half the number of targetable alleles as self-fertile hermaphrodites, and the oocyte allele could be edited in both germ cells and embryos. (A-H) Cas9-targetable polymorphic alleles of loci are shown in green, and non-targetable polymorphic alleles are shown in gray. The far-left column shows the configuration of oocyte and sperm alleles [maternal (m) or paternal (p)] for the *dpy*-*10* co-conversion marker and *sex*-*1* or *aft*-*2* in mated or selffertile hermaphrodites. The ovals display the genotypes of *sex*-*1* or *atf*-*2* loci in Dpy or Rol progeny of mated (blue) or self-fertile (red) hermaphrodites as determined by sequence analysis. An asterisk indicates the polymorphic allele with an indel. This allele is shown in parenthesis because the indel occasionally changes the allele-specific sequence. For self-fertile hermaphrodites in (B, E, and H) and for mated hermaphrodites in (A), either the sperm or oocyte allele could be repaired imprecisely to form an indel. Thus, only one arbitrary allele is shown with an indel. For (C-E) the average percentage of indels ± SEM is provided in blue (mated hermaphrodites) or red (self-fertile hermaphrodites), and the percentages for individual replicates are shown in black. The numbers in brackets below the percentage of indels show the fractions of total indel-laden progeny that have identical homozygous indels. For mated hermaphrodites, the percentage of *sex*-*1* or *aft*-*2* indels within Dpy or Rol outcrossed progeny (n) was calculated by the formula: (number of indels in target gene) / (total number of Rol or Dpy F1 outcrossed progeny) X 100. For self-fertile hermaphrodites, the percentage of *sex*-*1* or *aft*-*2* indels within total Dpy or Rol progeny (n) was calculated by the formula: (number of indels in target gene) / (total number of Rol or Dpy F1s in self progeny) X 100.

In subsequent experiments, we specifically targeted either the paternal allele (male sperm genome) or maternal allele (hermaphrodite oocyte genome) of a locus in mated hermaphrodites. In principle, because Cas9 has the opportunity to cleave hermaphrodite oocyte DNA both in developing germ cells and in embryos, the efficiency of editing maternal DNA might be expected to exceed the efficiency of editing paternal DNA, which must occur only after fertilization in these experiments. We found the opposite, consistent with the mating experiment above.

A Cas9-targetable polymorphic allele of a locus was introduced into the embryo from either the hermaphrodite oocyte or male sperm. This allele had a single nucleotide polymorphism in the penultimate position of the target-specific DNA sequence within the spacer seed sequence, permitting allele-specific Cas9 cleavage without off-target cleavage of wild-type alleles (Materials and Methods). For the gene *sex*-*1*, when the targetable polymorphic AG allele was contributed from the male sperm, an average of 88% (± 2) of *dpy*-*10* mutants had an indel in the paternal allele of *sex*-*1* (Figure 10C). In contrast, when the targetable AG allele was contributed from the hermaphrodite oocyte, an average of 43% (± 4) of *dpy*-*10* mutants had an indel in the maternal *sex*-*1* allele (Figure 10D) (*P* < 10^−5^, chi-square). The rate of mutagenesis for the paternal AG allele was also twice that for the identical AG alleles in homozygous self-fertile hermaphrodites (43% ± 6) (*P* < 10^−5^, chi-square), even though the male sperm genome had half the number of targetable alleles as self-fertile hermaphrodites, and hermaphrodite alleles could be edited in both germ cells and embryos, while the paternal allele could only be edited in the embryo (Figure 10E).

We conclude that (i) highly efficient genome editing does indeed occur in embryos; (ii) male sperm DNA can be targeted more effectively than hermaphrodite oocyte or sperm DNA; and (iii) the act of mating *per se* does not increase the efficiency of genome editing. Our discovery of homozygous identical indels in the progeny of self-fertile hermaphrodites supports the conclusion that genome editing is efficient in embryos after pronuclear fusion (Figure 10, B, E, and H). Homozygous indels arose from Cas9 cleavage of the unmutagenized parental genome in the embryo followed by HDR using a homologous chromosome with an indel for the template. Even when these homozygous indels (Figure 10E) were counted as independent Cas9 events and added to those causing the original heterozygous indels, the total frequency of Cas9-mediated repair in hermaphrodite sperm and oocyte genomes (59%) was less than the frequency in paternal genomes of cross-progeny embryos (88% *P* = 0.002, chi-square). Furthermore, a larger number of independent, distinct indels were formed in male sperm genomes than in hermaphrodite sperm and oocyte genomes, indicating that male sperm genomes were targeted more effectively than genomes of hermaphrodite gametes. In mouse embryos, editing of sperm genomes also occurred more efficiently than editing of egg genomes (Suzuki *et al*. 2014).

These findings with *sex*-*1* were confirmed at two other loci, *aft*-*2 II* (Figure 10, F-H) and an intergenic locus on chromosome I (Figure S5A). At *aft*-*2* for example, 90% of *dpy*-*10* mutants had indels in male sperm alleles compared to 35% of *dpy*-*10* mutants with indels in hermaphrodite oocyte alleles (*P* < 10^−5^, chi-square) or 31% of *dpy*-*10* mutants with indels in hermaphrodite sperm or oocyte alleles (*P* < 10^−5^, chi-square) (Figure 10, F-H). The findings were also reproduced in a second set of *sex*-*1* experiments in which the wild-type GG allele was targeted rather than the polymorphic AG allele (Figure S5B).

While highly elevated rates of editing were observed for male sperm DNA at inter-and intragenic sites on three different chromosomes, an exception occurred at the *lir*-*2* locus (Figure S5C). The paternal allele was targeted at half the rate (36%) as alleles in self-fertile hermaphrodites (72%), revealing that mating can improve editing rates at many, but not all, loci.

### Precise DSB repair via inter-homolog HDR occurs at high frequency in embryos

Use of polymorphic alleles to restrict editing to either the paternal or the maternal genome enabled us to make two further comparisons. We assessed the efficiency of recovering mutations in the target gene when both the co-conversion marker and the target gene were cleaved and repaired in the same parental genome (either maternal or paternal) or instead in different genomes (Figure 11). We also compared the frequency of precise DSB repair in embryos via HDR from the homologous chromosome (inter-homolog HDR) to the frequency of imprecise repair via indel formation (Figure 11). The design of experiments in Figure 11 allowed us to quantify bona fide inter-homolog HDR events in embryos. This class of repair events had been inferred from the recovery of identical homozygous indels in prior experiments (Figure 10, B, E, and H),

**Figure 11.**
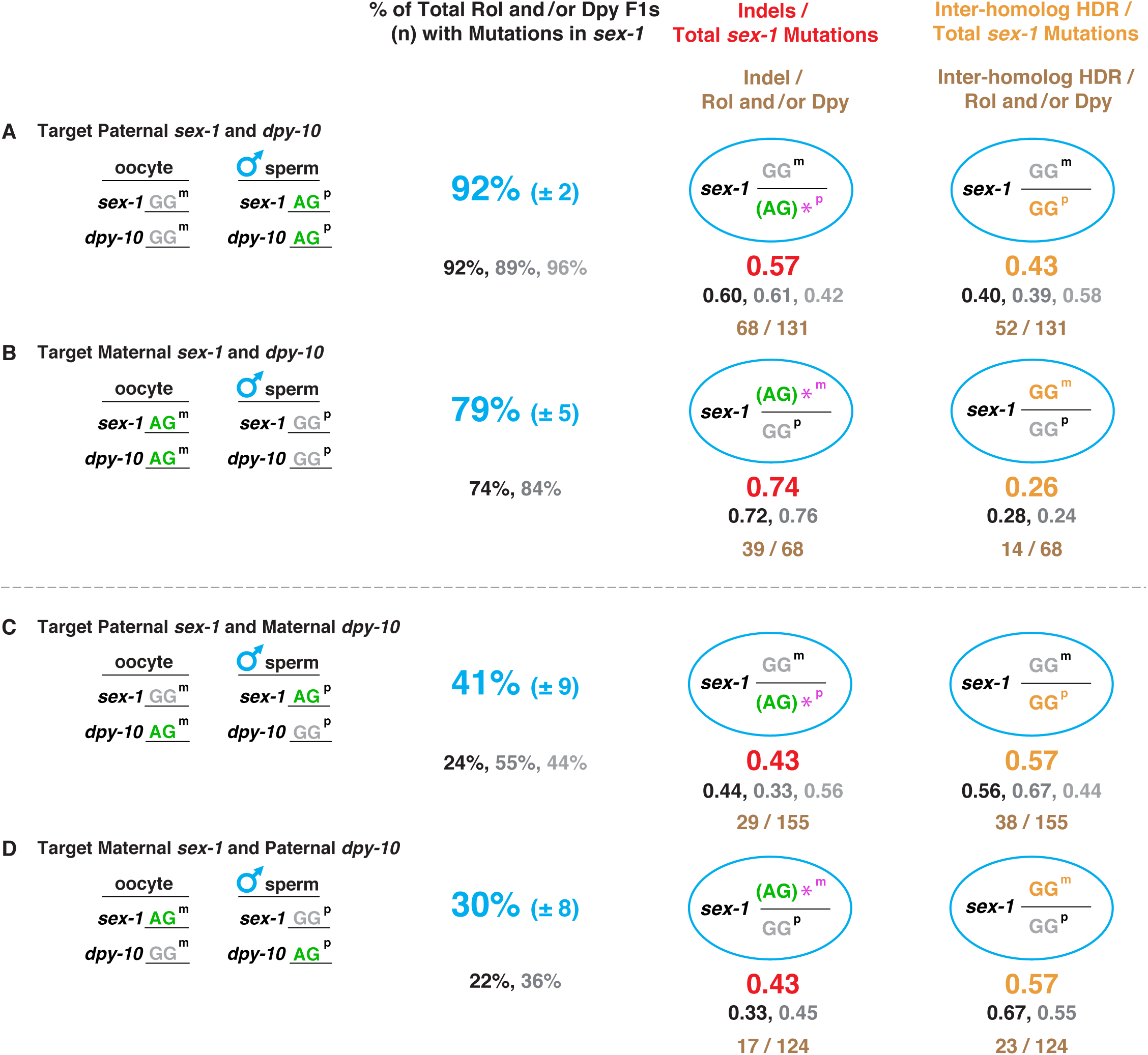
Efficiency of recovering genome editing events is greater if the co-conversion marker and target gene are cleaved and repaired in the same parental genome Comparison is shown of total *sex*-*1* mutation frequency (indels and inter-homolog HDR events) when the *dpy*-*10* co-conversion marker was targeted in the same (A, B) or in different (C, D) parental genomes (either paternal [p] or maternal [m]) as *sex*-*1*. Comparison is also shown for the frequency of imprecise DNA repair events via indel formation versus the frequency of precise DNA repair in embryos via inter-homolog HDR. Cas9-targetable polymorphic alleles of loci are shown in green, and non-targetable polymorphic alleles are shown in gray. The far-left column shows the configuration of hermaphrodite oocyte (m) and male sperm (p) alleles for *dpy*-*10* and *sex*-*1*. Frequency of total *sex*-*1* mutations relative to the total number (n) of Rol or Dpy cross progeny was calculated by the formula: (total number of *sex*-*1* indels and inter-homolog HDR events) / (total number of Rol or Dpy F1s) X 100. The average of percentages for all replicates of total *sex*-*1* mutations ± SEM are shown in blue, and the percentages for individual replicates are shown in black, medium gray, and light gray. Frequency in obtaining mutations in *sex*-*1* (combination of indels and inter-homolog HDR events) was invariably higher if both the marker and target gene were cleaved and repaired in the same parental genome, either paternal or maternal (*P* < 10^−5^, chi-square). The left column of ovals shows the genotypes of progeny with *sex*-*1* indels in Dpy or Rol outcrossed progeny, as determined by sequence analysis. An asterisk indicates the polymorphic allele with an indel. This allele is shown in parenthesis because the indel occasionally changes the AG sequence. The fraction of *sex*-*1* indels included in the total number of *sex*-*1* mutations (indels and inter-homolog HDR events) for all replicates combined is shown in red. The total number of indels compared to the total number of *sex*-*1* mutations for all replicates combined is shown in brown. The right column of ovals shows the genotypes of *sex*-*1* inter-homolog HDR events in Dpy or Rol outcrossed progeny, as determined by sequence analysis. The fraction of *sex*-*1* interhomolog HDR events included in the total number of *sex*-*1* mutations for all replicates combined is shown in orange. The respective replicates for fractions of indels and inter-homolog HDR events are shown black, medium gray, and light gray, matched to replicates for total percentages of *sex*-*1* mutations in each experiment. The total number of inter-homolog HDR events compared to the total number of *sex*-*1* mutations for all replicates combined is shown in brown. Remarkably, precise DSB repair via inter-homolog HDR, like indel formation, occurs at high frequency in embryos from mated hermaphrodites. In (C), quantification of inter-homolog events was based on the assumption that 100% of the progeny were cross-progeny. The validity of this assumption was based on our observation that all Dpy or Rol progeny in (A, B, and D) were cross-progeny, as determined by the criteria given in Materials and Methods.

For the first comparison, the success in obtaining precise or imprecise mutations in *sex*-*1* was invariably higher if both the marker and gene were targeted in the same parental genome, either paternal or maternal (*P* < 10^−5^, chi-square). When paternal alleles for both the marker and *sex*-*1* were targeted, 92% (± 2) of *dpy*-*10* mutants had a mutation (precise or imprecise) in the paternal *sex*-*1* allele (Figure 11A). In contrast, only 41% (± 9) of *dpy*-*10* mutants had a mutation in the paternal *sex*-*1* allele when the maternal *dpy*-*10* allele was targeted (*P* < 10^−4^, chi-square) (Figure 11C). Similarly, when maternal alleles were targeted for both the marker and *sex*-*1*, 79% (± 5) of *dpy*-*10* mutants had a mutation in the maternal *sex*-*1* allele (Figure 11B). However, only 30% (± 8) of *dpy*-*10* animals had a mutation in the maternal *sex*-*1* allele when the paternal *dpy*-*10* allele was targeted (*P* < 5 × 10^−4^, chi-square) (Figure 11D).

For the second comparison, a surprisingly high fraction of DSB repair events occurred precisely via inter-homolog HDR (0.26-0.57 of total *sex*-*1* mutations), when either the maternal or paternal *sex*-*1* locus was targeted for cleavage and repair (Figure 11, A-D). This result indicates that homology-directed repair occurs at high frequency in embryos and that the repair must have taken place after fusion of the paternal sperm and maternal oocyte pronuclei, when homologs were in proximity. Repair of Cas9 DSBs through inter-homolog HDR has also been observed in mice and tomatoes (Wu *et al*. 2013; Filler Hayut *et al*. 2017; Ma *et al*. 2017).

### Precise homology-directed repair from an exogenous template occurs at high frequency in embryos

The high frequency of inter-homolog HDR events in cross-progeny embryos (Figure 11) raised the question of whether the frequency of HDR using exogenous templates would also be high in cross-progeny embryos. We determined the frequency of HDR from exogenous templates and compared it directly with that of inter-homolog HDR and imprecise repair (Figure 12, A and B). When only paternal alleles of both *sex*-*1* and *dpy*-*10* were targeted in cross-progeny embryos, HDR events from exogenous templates occurred at a good frequency (24% of all *sex*-*1* mutations), and they were correlated with a reduction of both inter-homolog HDR events (reduced from 33% to 23%, *P* < 10^−3^, chi-square) and imprecise repair (reduced from 67% to 53%, *P* < 10^−5^, chi-square) (Figure 12A). Because paternal alleles can only undergo repair in embryos, and each form of repair precludes the others, all forms are predicted to be in direct competition, as observed in our data.

**Figure 12.**
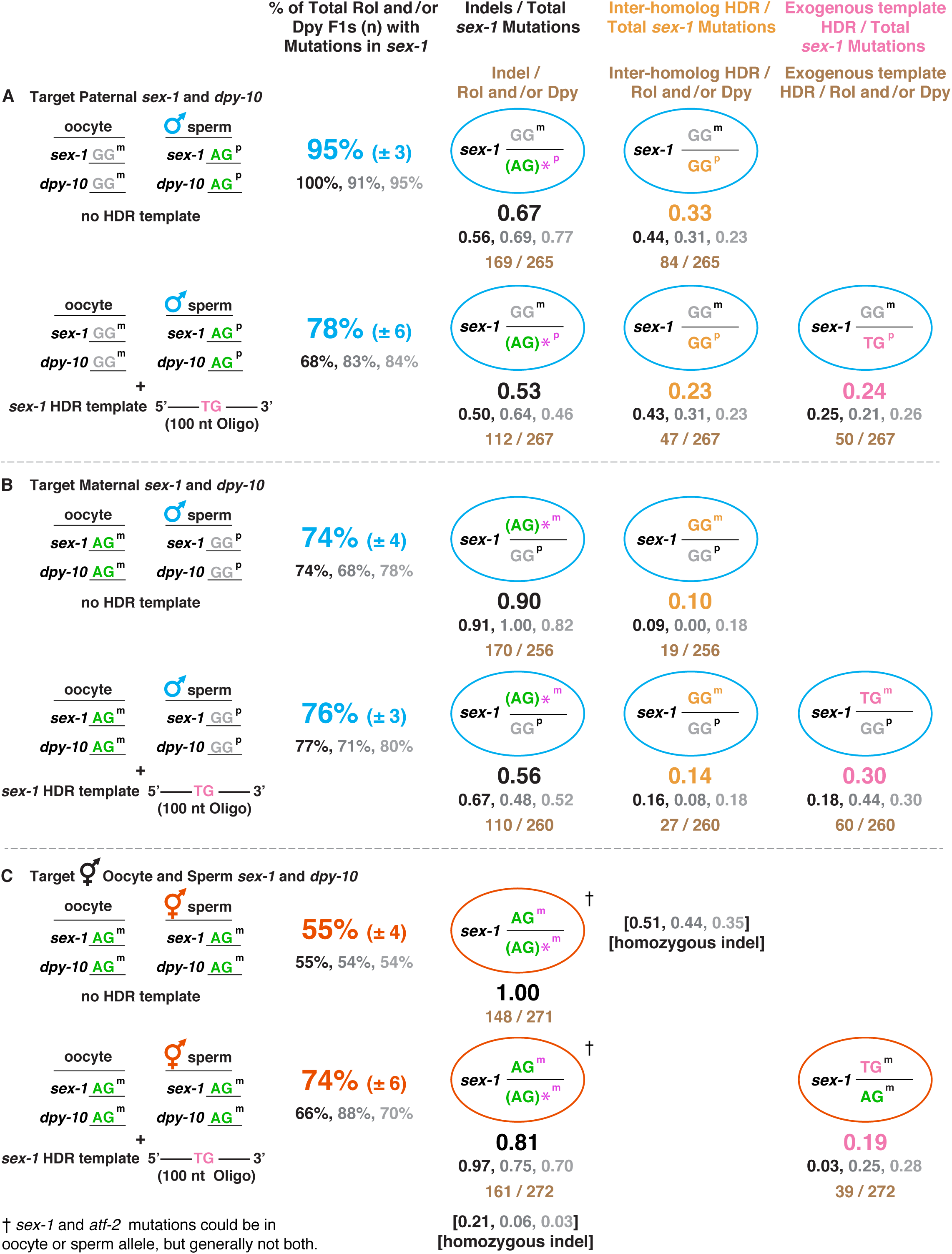
Homology-directed repair from an exogenous repair template occurs at high frequency in embryos (A,B) Comparison is shown for total *sex*-*1* mutation frequency (blue), including either indels and precise inter-homolog HDR events [top] or indels, precise inter-homolog HDR events, and precise HDR from an exogenous single-stranded template [bottom] when the *dpy*-*10* co-conversion marker was targeted in mated hermaphrodites in the same parental genomes as *sex*-*1*, either paternal [p] in (A) or maternal [m] in (B). The 100 nt single-stranded HDR repair template had only a single nucleotide change from the wild-type sequence. The AG sequence immediately 5′ of the PAM in the protospacer strand was changed to TG. Comparison is also shown for the fraction of indels (black, left column of ovals), the fraction of precise inter-homolog HDR events (orange, middle column of ovals), and the fraction of precise HDR events from the exogenous repair template (pink, right column of ovals). The replicates for each category are shown in black, medium gray, and light gray. The configurations of hermaphrodite oocyte (m) and male sperm (p) alleles for *dpy*-*10* and *sex*-*1* are displayed as in the left column of Figure 11, A and B. Genotypes of progeny with indels and inter-homolog HDR events are displayed like the right and left ovals of Figure 11, A and B. The formula for calculations and method of displaying replicates is the same as in Figure 11, A and B. The right column of ovals shows the genotypes of progeny with precise HDR from the exogenous repair template, as determined by sequence analysis. Shown in brown for each column of ovals is either the total number of *sex*-*1* indels found in all Rol and/or Dpy progeny from all replicates, the total number of *sex*-*1* inter-homology HDR events found in all Rol and/or Dpy progeny from all replicates, or the total number of *sex*-*1* HDR events via exogenous templates found in all Rol and/or Dpy progeny from all replicates. These experiments reveal that HDR using exogenous templates occurs in cross-progeny embryos from male/hermaphrodite matings when either the maternal (oocyte) or paternal (sperm) allele is targeted for Cas9-mediated DNA cleavage. (C) Comparison is shown among progeny of self-fertile hermaphrodites of the frequency (red) of imprecise DNA repair via indel formation (top) versus the frequency (red) of imprecise DNA repair (indels) and precise DNA repair via HDR from the exogenous 100 nt single-stranded template [bottom]. The genotypes of parents and progeny are displayed as in Figure 10E. Experimental replicates showing the fraction of indels (top) and fraction of indels and HDR events (bottom) are displayed in black, medium gray and light gray. Numbers in brackets represent the fraction of total indel-laden progeny that have identical homozygous indels. Shown in brown is the total number of *sex*-*1* indels or the total number of *sex*-*1* HDR events from exogenous templates found in all Rol and/or Dpy mutants from all combined replicates. The total frequency of Cas9 editing events reported in the text, when taking into account homozygous identical indels as independent Cas9 cleavage events, was calculated by the formula: [(fraction of total Rol and/or Dpy F1s with mutations in *sex*-*1*) + (fraction of total Rol and/or Dpy F1s with homozygous identical indels in *sex*-*1*)] X 100. These experiments reveal that genomes of embryos from self-fertile hermaphrodites undergo a substantial number of HDR events from exogenous templates.

When only maternal alleles of both *sex*-*1* and *dpy*-*10* were targeted in cross-progeny embryos, HDR events from exogenous templates also occurred at good frequency (30% of all *sex*-*1* mutations), one not statistically different from related HDR events at paternal alleles (*P* = 0.2) (Figure 12B). However, in contrast to HDR events with exogenous templates at paternal alleles, these HDR events at maternal alleles correlated with a loss of imprecise repair (reduced from 90% to 56%, *P* < 10^−5^, chi-square), but not a loss of inter-homolog HDR (changed from 10% to 14%, *P* = 0.2, chi-square). These results suggest that for maternal alleles, imprecise repair and HDR using exogenous templates occur in a similar window of time, during meiotic prophase or until pronuclear fusion in embryos, but interhomolog HDR occurs later. Both the insignificant change in frequency of inter-homolog HDR events when exogenous HDR repair templates were present and the general low frequency of inter-homolog
HDR events are consistent with inter-homolog HDR occurring after pronuclear fusion in embryos, once other forms of repair had occurred.

We also asked whether HDR from exogenous templates occurs in embryos of self-fertile hermaphrodites. We found that 19% of total *sex*-*1* mutants had an HDR event from an exogenous template (Figure 12C bottom). Moreover, the total frequency *dpy*-*10* co-converted animals having a *sex*-*1* mutation rose from 55% (± 4), when only heterozygous indels were scored in experiments lacking a homologous repair template (Figure 12C top), to 74% (± 6) when both indels and HDR events were scored in experiments with a homologous repair template (*P* < 10^−5^, chi-square) (Figure 12C bottom). This latter result revealed that quantifying only heterozygous indels in progeny of self-fertile hermaphrodites did not reflect the total frequency of independent editing events. Indeed, inclusion of inter-homolog HDR events deduced from the occurrence of homozygous identical indels in experiments lacking an exogenous repair template increased the total frequency of *sex*-*1* editing events from 55% to 78%, (see calculation in Figure 12C legend), a frequency not different from the 74% found when including HDR events from exogenous templates.

To infer where HDR via exogenous templates had occurred, we compared the proportion of heterozygous indels and homozygous indels in progeny of self-fertile hermaphrodites that had been given Cas9 RNPs either with or without an exogenous HDR template. We found that HDR repair from an exogenous template correlated with a reduction in fraction of homozygous identical indels (*P* < 10^−5^, chi-square) but not a reduction in the fraction of unique heterozygous indels (*P* = 0.3, chi-square). Thus, HDR from an exogenous template appeared to have competed with inter-homolog HDR, which can only occur in embryos after fusion of oocyte and sperm pronuclei. It did not compete with imprecise repair, which can occur during meiotic prophase or in embryos. We infer that genomes in embryos from self-fertile hermaphrodites undergo a substantial number of HDR events from exogenous templates, most likely after pronuclear fusion.

## Conclusions

Our Cas9 editing strategies have achieved highly efficient genome engineering in *C. elegans*. These innovations can be exploited to enhance genome editing in diverse species.

First, our detailed analysis of imprecise repair outcomes in *C. elegans*, along with data from other species (Mali *et al*. 2013b; Van Overbeek *et al*. 2016; Richardson *et al*. 2018), showed that non-templated repair at Cas9 cleavage sites is asymmetric, favoring deletions and insertions 5′ of the PAM on the protospacer strand. Thus, imprecise repair can be increased significantly by selecting Cas9 targets with PAMs located 3′ of desired changes.

Second, our systematic analysis also showed that HDR from a single-stranded repair template is directional and consistent with a synthesis-dependent strand annealing mechanism. We exploited this finding to devise guidelines for inserting precise changes with high frequency into sites within close proximity (30 bp) to the DSB. Insertions 5′ of the PAM can be achieved efficiently using a single-stranded repair template corresponding to the protospacer. In contrast, precise changes 3′ of the PAM are achieved most reliably using a spacer strand repair template instead. Both spacer and protospacer strand templates are effective for inserting precise changes directly at the DSB site. Because the bound Cas9 nickase RuvC also has a 3′ - 5′ exonucleolytic activity that catalyzes limited resection of the cleaved protospacer strand, a spacer strand repair template can elicit not only high-efficiency insertion of polymorphisms 3′ of the PAM but also low-efficiency insertion of polymorphisms just 5′ of the PAM.

Third, we overcame limitations related to inserting long non-homologous fragments of DNA near a DSB and to engineering small specific changes at considerable distance from a DSB. These desired outcomes had been problematic because HDR from a double-stranded repair template is inefficient, distance-dependent, and highly directional, favoring changes 5′ of the PAM. The strategy that surmounted these limitations uses two different Cas9 guides to create two DSBs that flank the region intended for insertion of exogenous DNA. This approach is effective for inserting ~ 10 kb of non-homologous DNA and for incorporating a series of nucleotide substitutions along the entire length of a region, up to 1.5 kb from a DSB. The scheme is also beneficial for editing insertion sites such as AT-rich introns that are not adjacent to DNA sequences necessary for effective Cas9 guide design.

Use of two Cas9 targets required that we determine the orientation of PAMs most successful for editing. One orientation causes the PAM to remain on the chromosome end after DNA excision; the other causes the PAM to reside on the DNA sequence excised from the chromosome. Editing is most efficient if both PAMs will remain on chromosome ends after DNA excision.

Fourth, we optimized Cas9 delivery methods and guide RNA design. We assessed the relative effectiveness of using either pre-assembled ribonucleoprotein complexes (RNPs) of guide RNAs and Cas9 or DNA expression vectors to deliver editing reagents to gonads. Using both delivery methods, we also assessed the effectiveness of guide RNAs differing only in the penultimate nucleotide at the 3′ end of their target-specific sequences (GG, AG, CG or UG). We found that Cas9 editing is more robust using RNPs than using DNA expression vectors. GG guides are crucial for successful editing using DNA expression vectors and are the most effective guides when using RNPs.

Fifth, we expanded the repertoire of easily scorable co-conversion markers used to identify Cas9-edited animals that likely also carry edits in targets of interest. The markers dramatically narrow the search for edited loci that fail to cause visible phenotypes. The *C. elegans* marker *zen*-*4* is selectable, causing edited animals to thrive in a population of temperature-sensitive lethal mutants. The second marker *ben*-*1* serves as a co-conversion marker appropriate for diverse nematode species. Cas9-induced *ben*-*1* mutations confer dominant resistance to the drug benomyl in both *C. elegans* and *C. briggsae*, allowing animals of both species to be mobile rather than paralyzed in the presence of the drug.

Sixth, we explored the timing, location, frequency, sex-dependence, and categories of DSB repair events. For this purpose, we designed allele-specific targets of Cas9 to be contributed during mating from either male or hermaphrodite germ cells. We found that male sperm DNA is generally more permissive to Cas9 editing than DNA from hermaphrodite germ cells. Furthermore, the frequency of recovering repair events in the target gene of interest is higher if editable alleles of both the target gene and co-conversion marker are introduced from the same parent during mating. Lastly, both imprecise repair and homology-directed repair from either exogenous repair templates or homologous chromosomes occur at unexpectedly high frequencies after fertilization in embryos.

## Acknowledgments

We thank T. Cline, A. Villeneuve, and members of the Meyer lab for insightful discussions. Some strains in this study were provided by the *Caenorhabditis* Genetics Center, which is funded by the National Institutes of Health (NIH) Office of Research Infrastructure Programs (P40 OD 010440). A.F.S. was supported by NIH grant R15GM117548 and the Center for Gene Regulation in Health and Disease. B.J.M. was supported in part by NIH grant R01GM030702. B.J.M. is an investigator of the Howard Hughes Medical Institute.

## Supplementary Figure Legends

**Figure S1.**
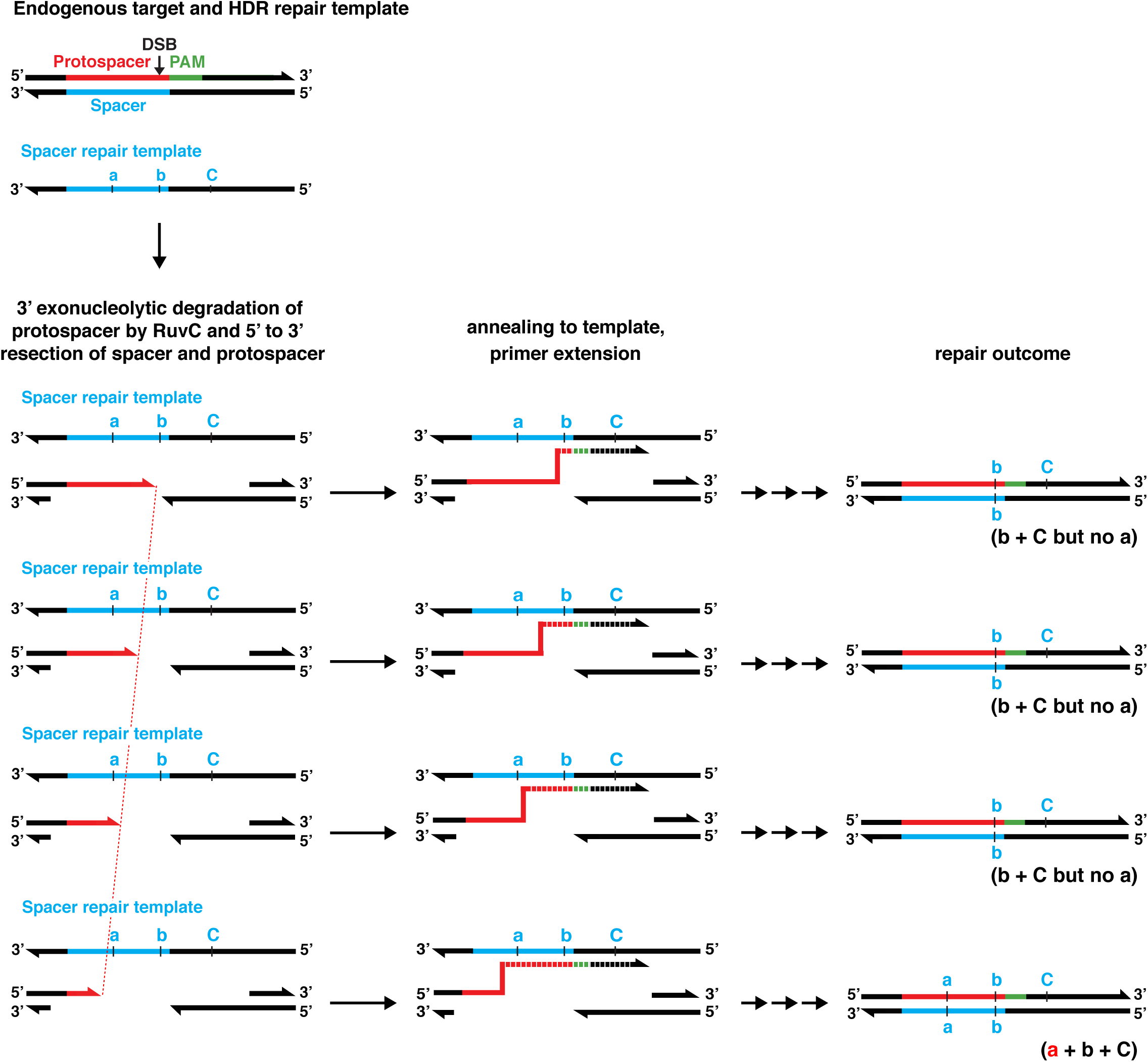
RuvC single-stranded 3′ exonucleolytic activity provides a plausible mechanism for insertion of polymorphisms 5′ of the PAM from a spacer strand repair template This model for the infrequent insertion of SNPs 5′ of the PAM from a spacer strand repair template requires the 3′ exonucleolytic activity of the single-stranded Cas9 nickase RuvC. This nickase resects single-stranded DNA *in vitro* up to 10 nucleotides in the 3′ - 5′ direction. Since polymorphism a is only 10 bp from the DSB, the RuvC nickase should occasionally resect the protospacer strand of the endogenous DNA past the location of the polymorphism and allow incorporation of this polymorphism from primer extension off the spacer repair template. The model shows different degrees of 3′ resection by RuvC on the protospacer strand (dashed red line) and the subsequent difference in repair outcomes for the inclusion of polymorphism a.

**Figure S2.**
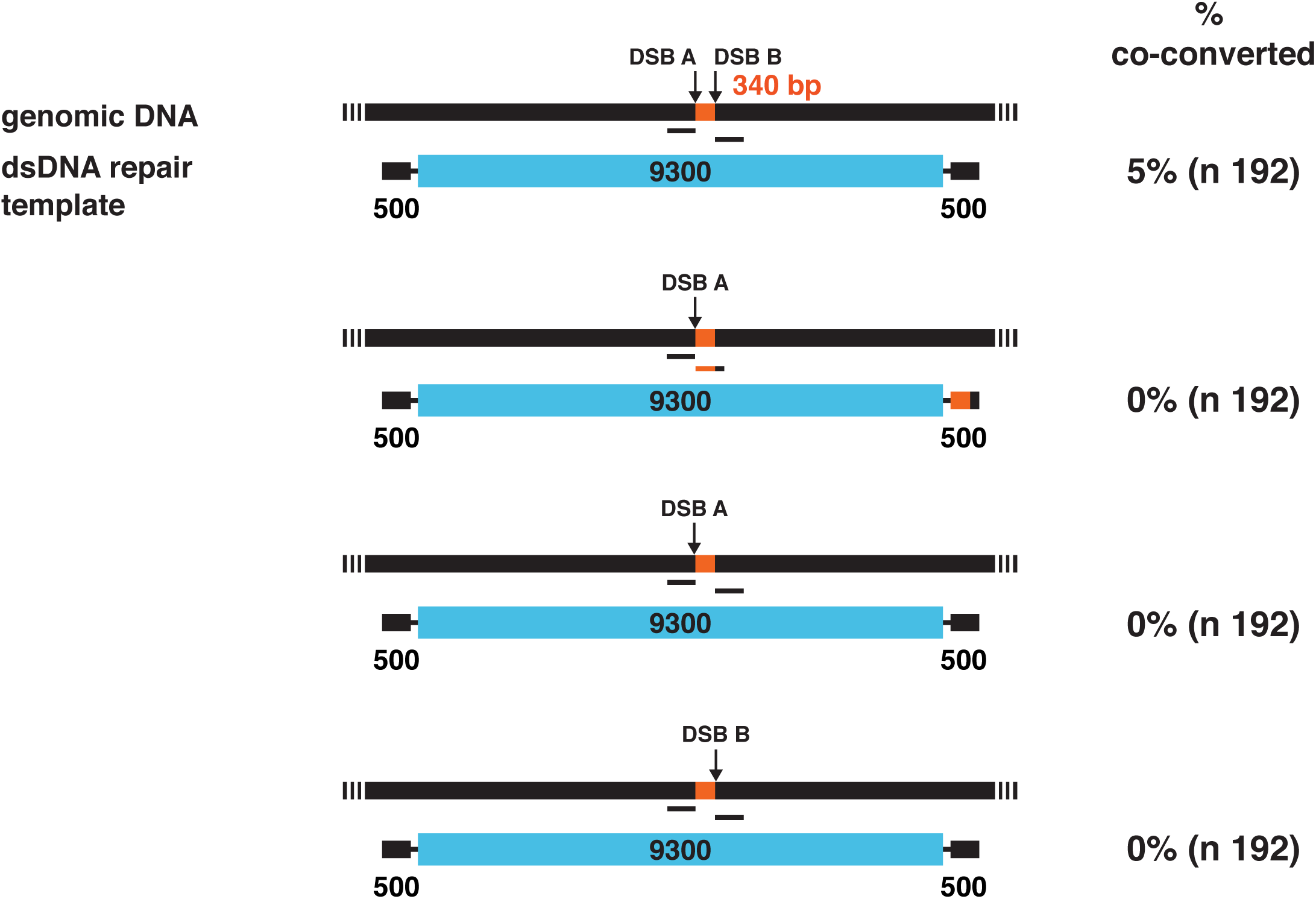
Insertion of large DNA fragments via double-stranded templates is infrequent at a single Cas9 cleavage site The strategy of using two guide RNAs and hence two DSBs 340 bp apart (orange bar) to insert large (9300 bp) fluorescent reporter transgenes into an endogenous site using a plasmid-based doublestranded repair template with 500 bp homology arms was successful in experiments with *dpy*-*10* as the co-conversion marker. See also Figure 5. However, co-conversion experiments were unsuccessful for inserting large stretches of DNA if they involved only one of the two guide RNAs and three different sets of 500 bp homology arms, as represented by the black and orange-black lines below the genomic DNA diagram.

**Figure S3.**
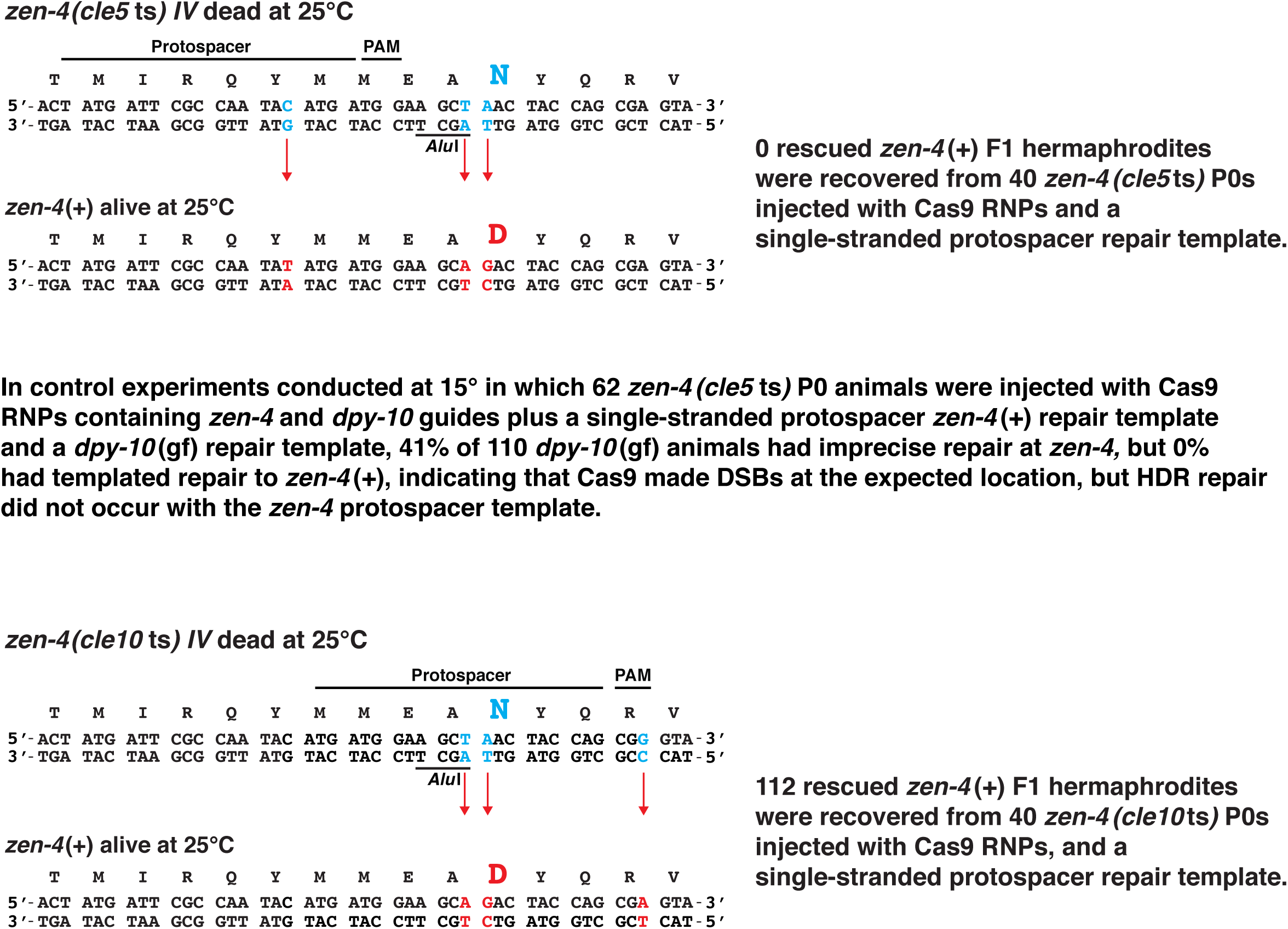
Strategy to develop a co-conversion marker that enabled selection of viable, edited animals (A) To develop *zen*-*4*(+) as a co-conversion marker we used Cas9 editing to create a temperature-sensitive lethal allele of *zen*-*4* that had the death-inducing GAC to AAC transition at codon 520 [*zen*-*4(cle5*ts)]. The strain also included an *Alu*I site that was made by converting codon 519 from a GCA alanine codon to a GCT alanine codon to distinguish edited genes from true revertants. These DNA changes were located 3′ of the PAM. In initial attempts to convert the *zen*-*4(cle5*ts) allele to a wild-type allele and thereby rescue the lethal phenotype, we used a protospacer stand repair template in which codon 515 had been converted to TAT from TAC to prevent the repaired *zen*-*4*(+) gene from being cleaved by Cas9. HDR failed, but imprecise repair succeeded, indicating that Cas9 RNPs cleaved the DNA at the expected location. This failure contributed to the evidence that protospacer strand HDR templates are not successful for repair of mutations 3′ of the PAM (see Figures 2 and 3). (B) We then created a second *zen*-*4* mutant strain (allele *cle10*ts) to change the location of Cas9 RNP binding. The strain carried the GAC to ACC transition at codon 520 to create the temperature-sensitive lethal mutation and also included the CGA to CGG transition at codon 523 to create a PAM near the target sequence for the guide RNA. In addition, an *Alu*I site was introduced by converting codon 519 from a GCA alanine codon to a GCT alanine codon to distinguish edited animals from true revertants. All these changes were 5′ of the PAM. Fortuitously, we chose a single-stranded HDR repair template that corresponded to the protospacer strand and found that the combination of guide and repair template was successful for HDR. This result reinforced the evidence in Figures 2 and 3 that a protospacer strand repair template is efficient for repair of sequences 5′ of the PAM.

**Figure S4.**
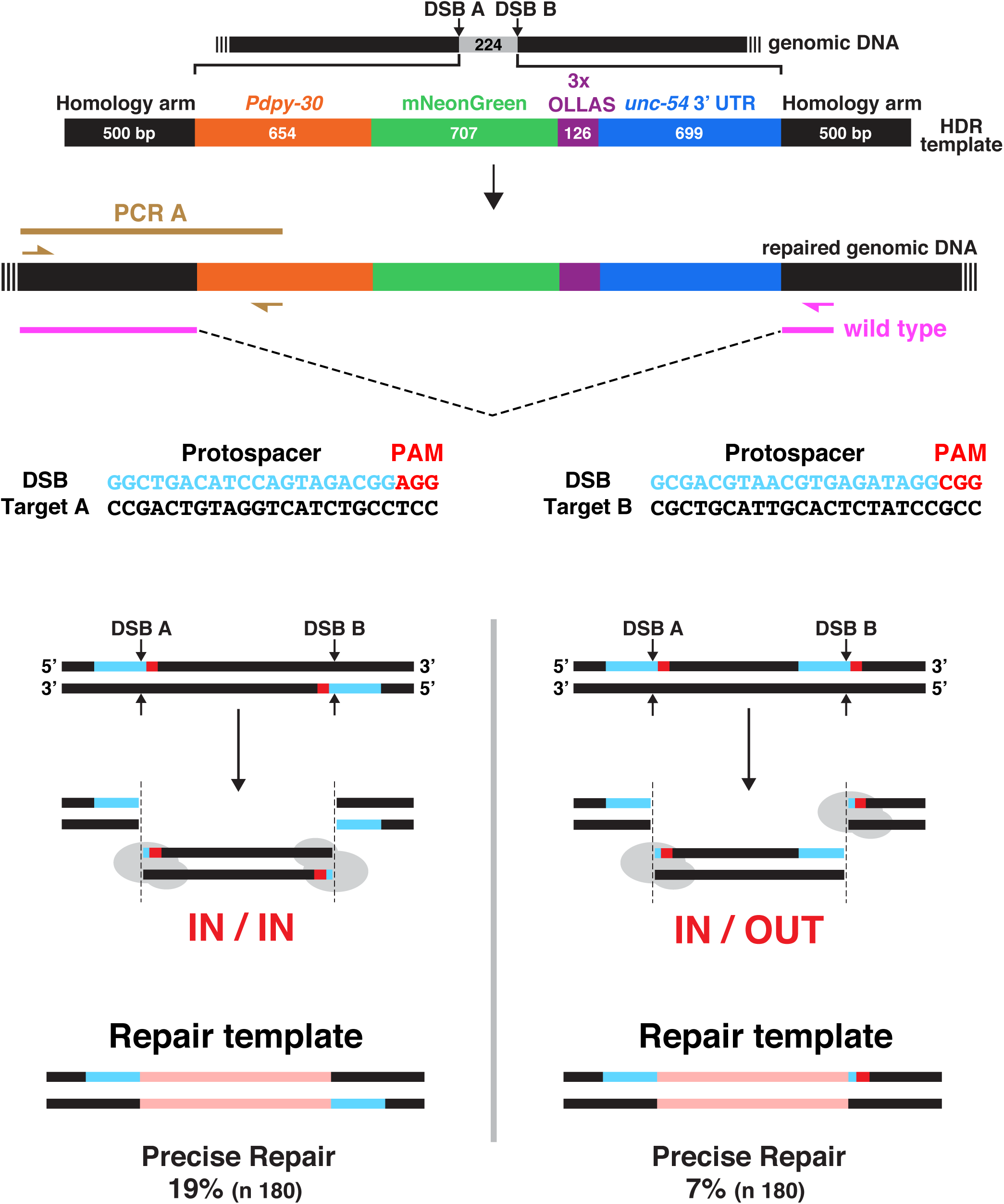
Orientation of PAMs at two nearby Cas9 cleavage sites influences HDR efficiency Cas9 was directed to two locations in close proximity (224 bp,Target A and Target B), to make two DSBs and thereby stimulate insertion of an exogenously supplied homologous repair template. To examine the effect of PAM orientation on HDR efficiency, two strains were made that had identical sequences at the insertion site except for the orientation of Target B. One strain had PAMs (red) in an IN/IN orientation, placing the PAMs in the excised genomic DNA, while the other strain had PAMs in an IN/OUT orientation, placing one PAM in the excised genomic DNA and the other at a free end of the cleaved chromosome. The mNeonGreen, double-stranded DNA reporter repair template had 500 bp homology arms corresponding to genomic sequences adjacent to the Cas9 DSBs. Repair templates with either the IN/IN or IN/OUT orientation were identical except for sequences corresponding to Target B sites. The IN/IN repair template included cleavage remnants of Target B that lacked a PAM, while the IN/OUT repair template included cleavage remnants of Target B that had the PAM (see diagram; reporter insert shown in pink). PCR was used to screen 180 Dpy or Rol F1s for each set of PAM orientations. A primer (left brown arrow) that annealed to genomic sequence located outside the bounds of the repair template and a second primer (right brown arrow) that annealed to the *dpy*-*30* promoter within the repair template only amplified sequences from worms having the desired genomic insertions at Target A. A PCR reaction using the primer (left brown arrow) outside the repair template and a third primer (pink arrow) within the right homology arm produced a 320 bp product from wild-type unedited genomic DNA. Percent (%) of Dpy or Rol F1s with precise insertions for each set of PAMs is shown. Homozygous insert-containing offspring for two positive F1s from each PAM orientation were subjected to Sanger sequencing. In all four cases, the edited genomes had precisely inserted repair templates. The percent of repair from the IN/IN orientation was statistically greater than that for the IN/OUT orientation (*P* = 0.001, chi-square)

**Figure S5.**
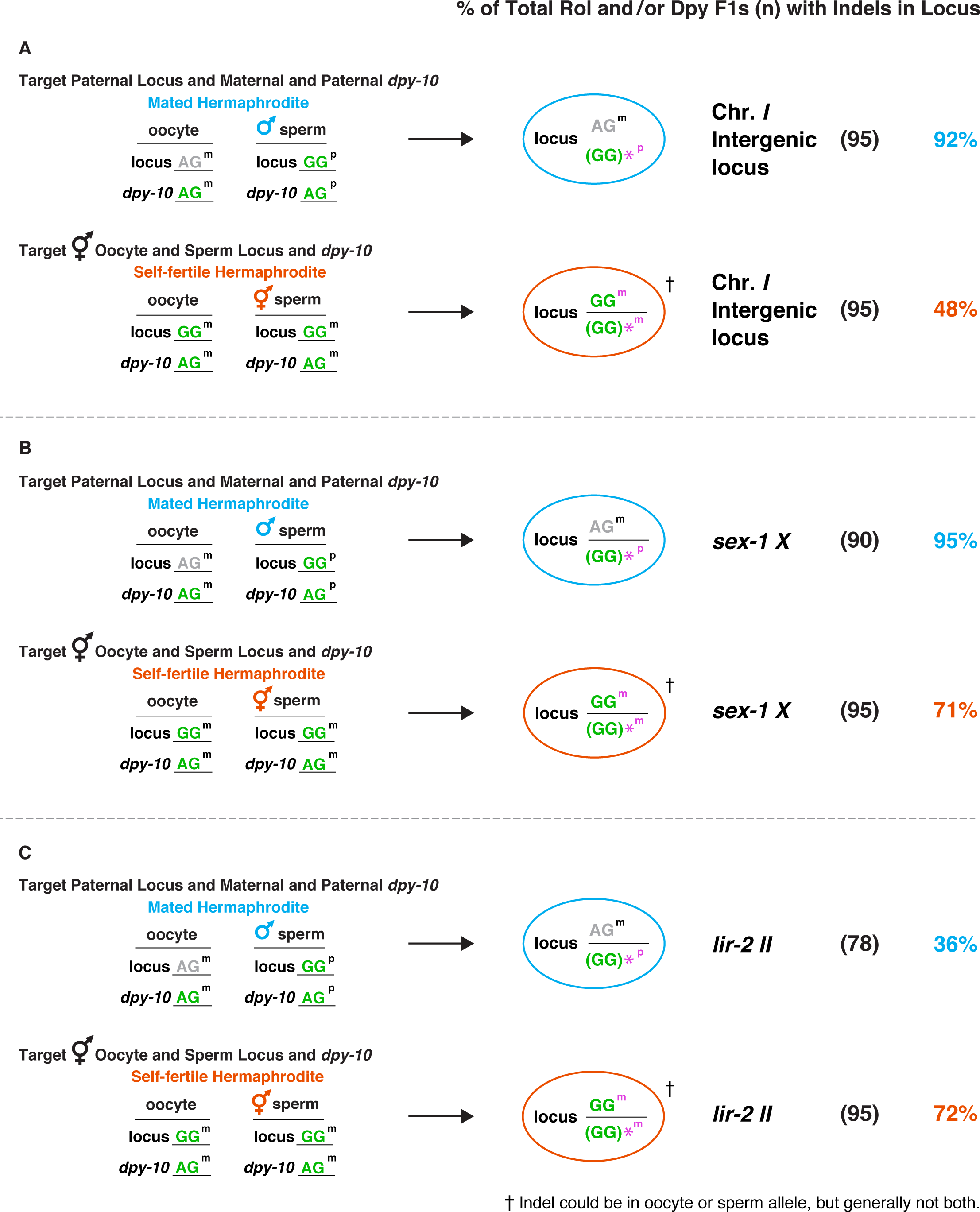
Genome editing occurs in embryos of mated hermaphrodites Comparison is shown for the frequency of indels at an intergenic locus on chromosome I (A), at *sex*-*1 X* (B) using a GG guide RNA instead of the AG guide in Figure 10, and at *lir*-*1 II* (C) in paternal (male sperm) genomes of progeny from mated hermaphrodites versus the frequency in sperm and/or oocyte genomes of self-fertile hermaphrodites. Cas9-targetable alleles of loci are shown in green, and non-targetable polymorphic alleles are shown in gray. The far-left column shows the configuration of oocyte and sperm alleles [maternal (m) or paternal (p)] for the *dpy*-*10* co-conversion marker and the designated locus in mated or self-fertile hermaphrodites. The ovals show the genotypes of designated loci in Dpy / Rol progeny from mated (blue) or self-fertile (red) hermaphrodites as determined by sequence analysis. An asterisk indicates the polymorphic allele with an indel. This allele is shown in parenthesis because the indel occasionally changes the GG sequence. For self-fertile hermaphrodites, either the sperm or oocyte allele could be repaired imprecisely to form an indel. Thus, only one arbitrary allele is shown with an indel. For mated hermaphrodites, the percentage of indels in Dpy / Rol outcrossed progeny (n) was calculated by the formula: (number of indels in target gene) / (total number of Rol / Dpy F1s in outcrossed progeny) X 100. For self-fertile hermaphrodites, the percentage of indels within Dpy / Rol progeny (n) was calculated by the formula: (number of indels in target gene) / (total number of Rol / Dpy F1s in self progeny) X 100. For *sex*-*1* and the intergenic locus on chromosome I, the frequency of indels in the single paternal allele of cross progeny from mated hermaphrodites was higher than that for the two identical alleles in progeny from self-fertile hermaphrodites (*P* < 10^−4^, chi-square), indicating that highly efficient genome editing occurs in embryos, and male sperm DNA can be targeted more effectively than hermaphrodite DNA. The frequency of indels at *sex*-*1* from self-fertile hermaphrodites is likely higher than that in Figure 10E, because GG guides are more effective than AG guides (see Figure 7). In contrast, the single paternal allele of *lir*-*2* was targeted at half the frequency as the two alleles in self-fertile hermaphrodites (*P* < 10^−5^, chi-square), indicating that mating does not always improve the frequency of mutagenesis.

**Table S1.**
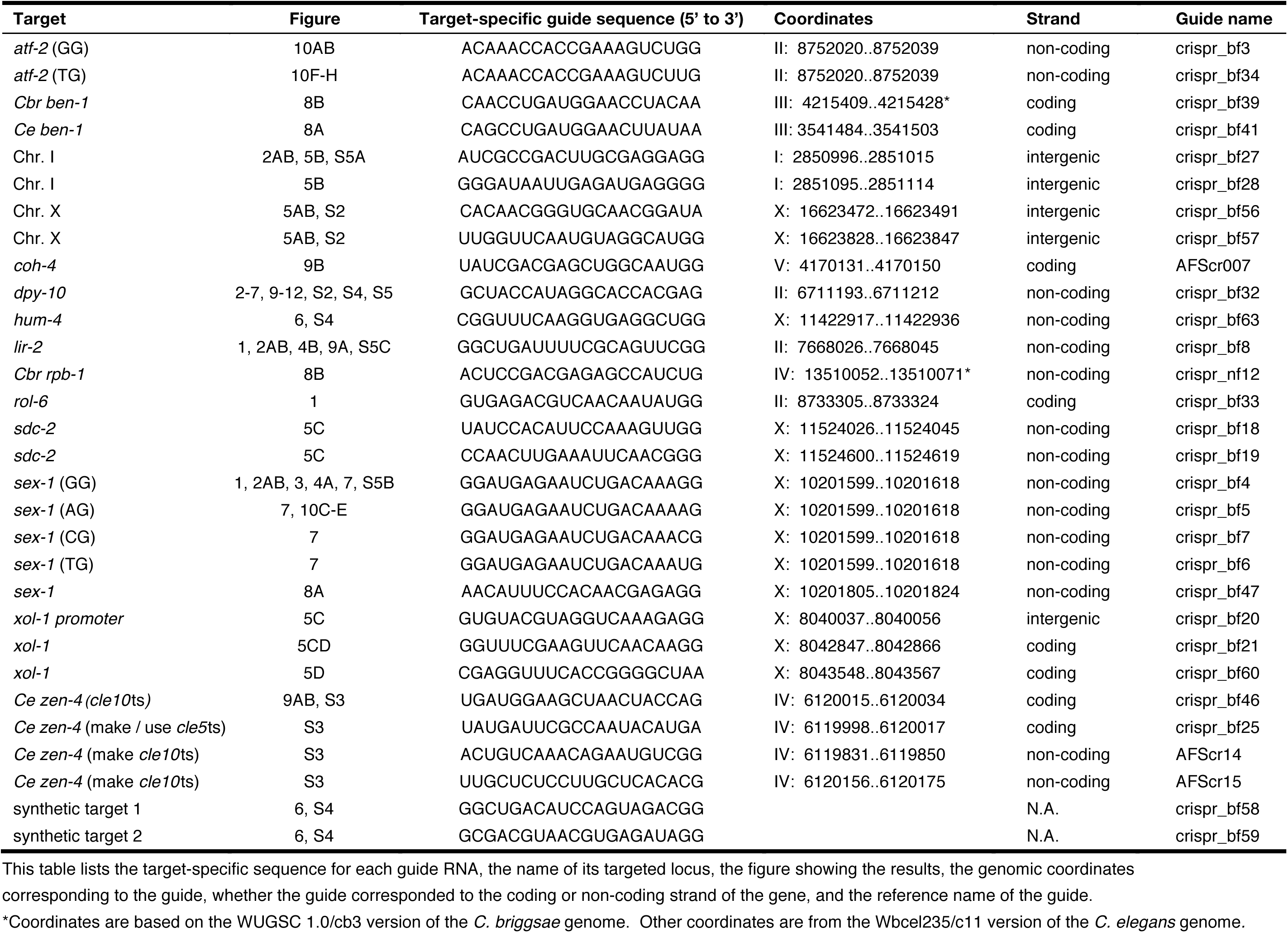
List of target-specific guide sequences.

**Table S2.**
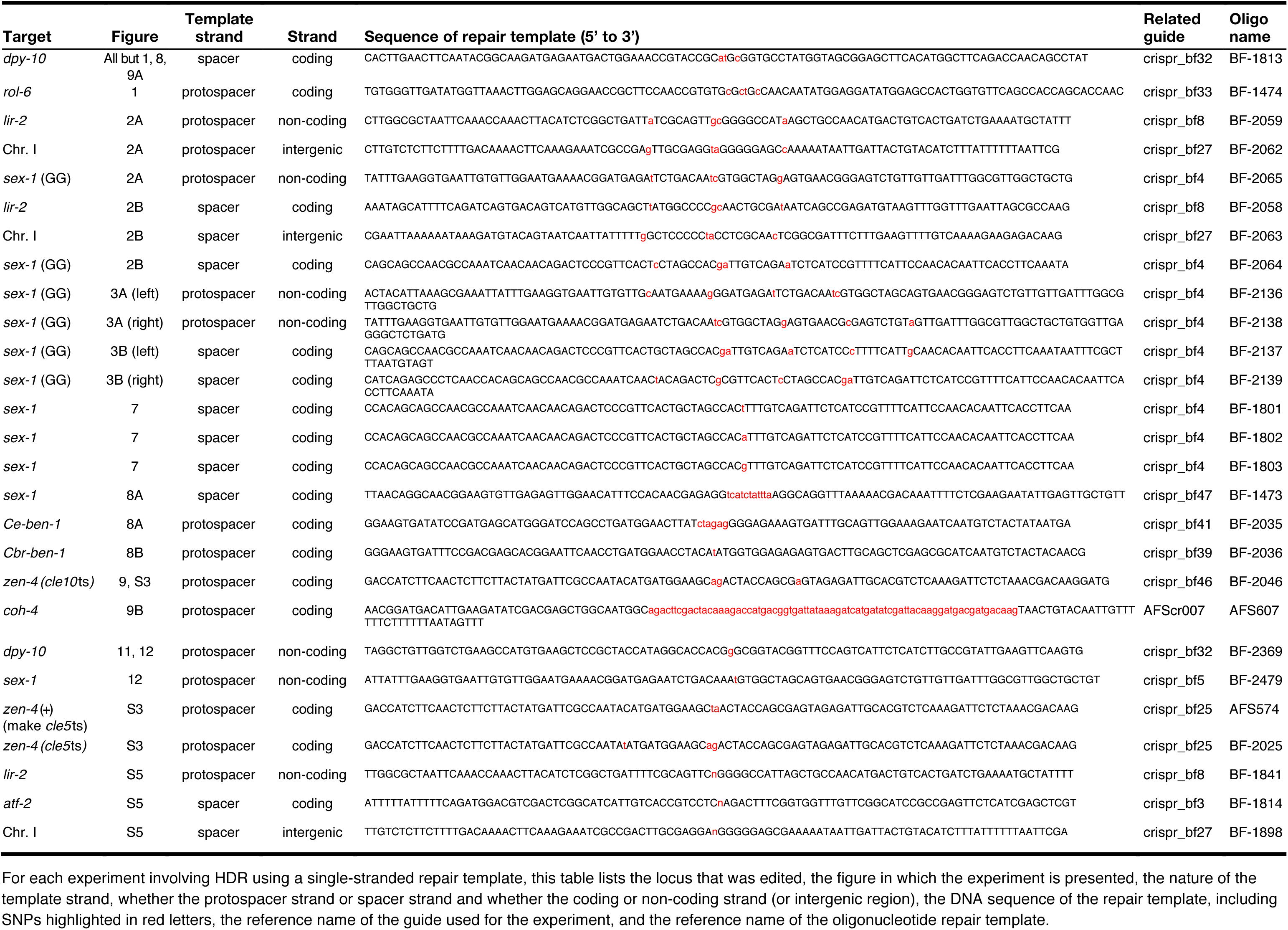
List of repair templates.

**Table S3.**
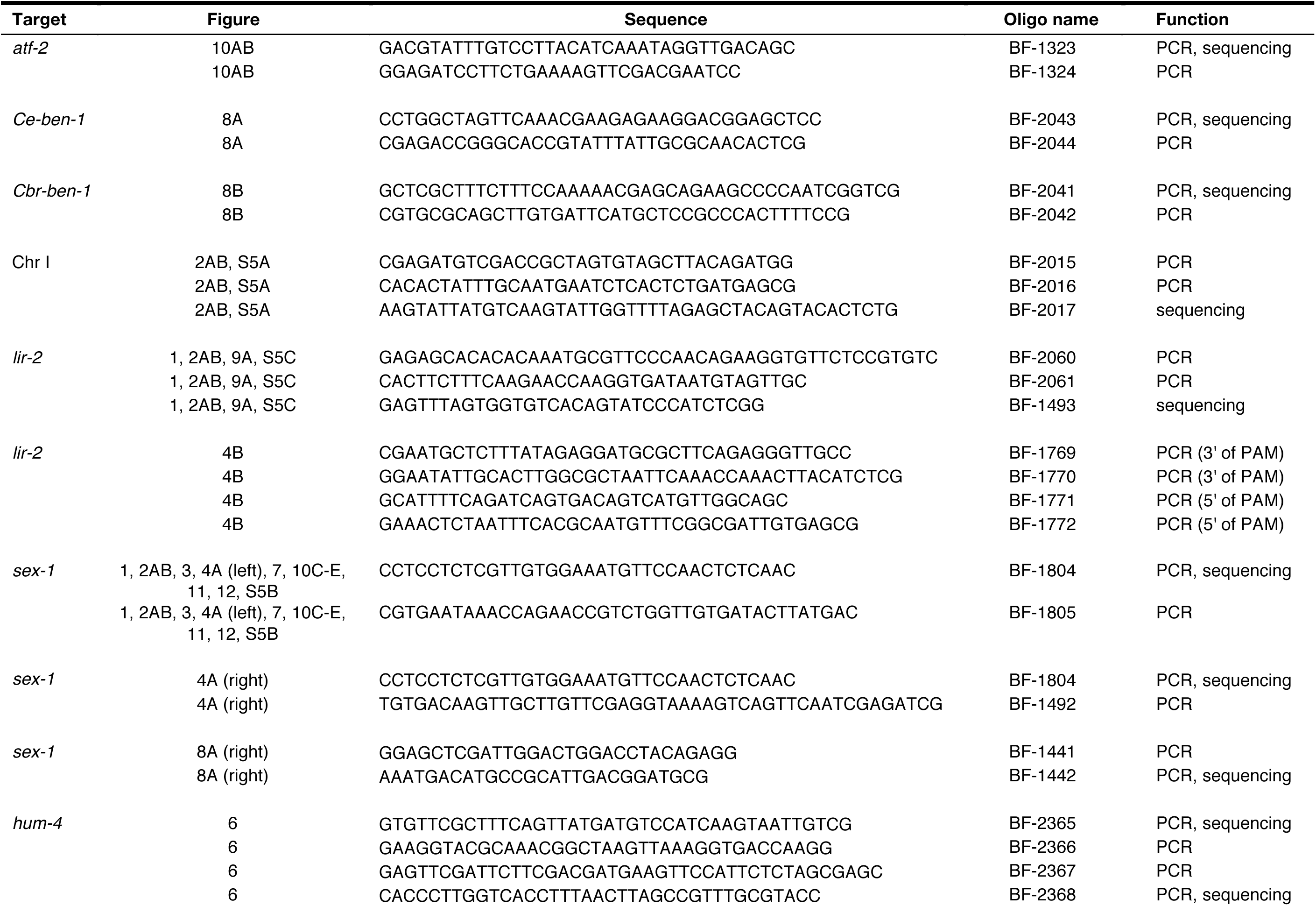
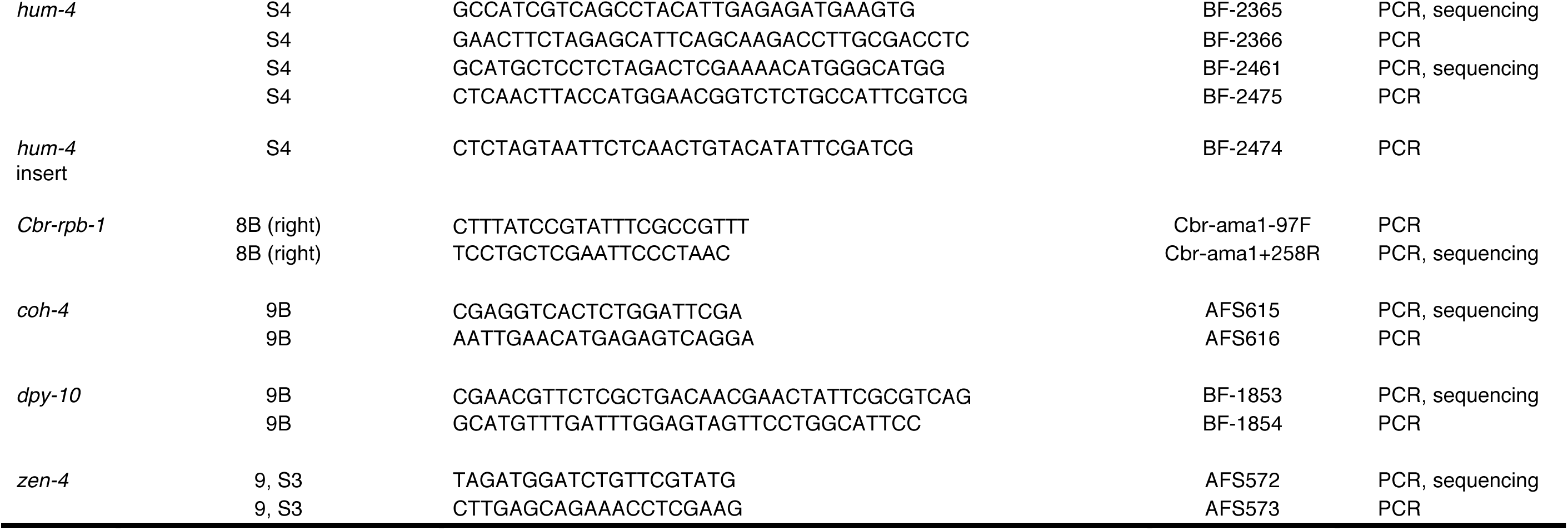
List of oligos for mutant sequencing.

